# Nach is a novel ancestral subfamily ofthe CNC-bZIP transcription factors selected during evolution from the marine bacteria to human

**DOI:** 10.1101/287755

**Authors:** Yuping Zhu, Meng Wang, Yuancai Xiang, Lu Qiu, Shaofan Hu, Zhengwen Zhang, Peter Mattjus, Yiguo Zhang

**Affiliations:** The Laboratory of Cell Biochemistry and Topogenetic Regulation, College of Bioengineering and Faculty of Sciences, Chongqing University, No. 174 Shazheng Street, Shapingba District, Chongqing 400044, China; Institute of Neuroscience and Psychology, School of Life Sciences, University of Glasgow, 42 Western Common Road, G22 5PQ, Glasgow, Scotland, United Kingdom; Department of Biochemistry, Faculty of Science and Engineering, Åbo Akademi University, Artillerigatan 6A, III, BioCity, FI-20520 Turku, Finland

**Keywords:** Nach, CNC, bZIP transcription factors, interaction network, evolution, transmembrane

## Abstract

All living organisms have undergone the evolutionary selection under the changing natural environments to survive as diverse life forms. All life processes including normal homeostatic development and growth into organismic bodies with distinct cellular identifications, as well as their adaptive responses to various intracellular and environmental stresses, are tightly controlled by signaling of transcriptional networks towards regulation of cognate genes by many different transcription factors. Amongst them, one of the most conserved is the basic-region leucine zipper (bZIP) family. They play vital roles essential for cell proliferation, differentiation and maintenance in complex multicellular organisms. Notably, an unresolved divergence on the evolution of bZIP proteins is addressed here. By a combination of bioinformatics with genomics and molecular biology, we have demonstrated that two of the most ancestral family members classified into BATF and Jun subgroups are originated from viruses, albeit expansion and diversification of the bZIP superfamily occur in different vertebrates. Interestingly, a specific ancestral subfamily of bZIP proteins is identified and also designated Nach (<u>*N*</u>rf <u>*a*</u>nd <u>*C*</u>NC <u>*h*</u>omology) on account of their highly conservativity with NF-E2 p45 subunit-related factors Nrf1/2. Further experimental evidence reveals that Nach1/2 from the marine bacteria exerts distinctive functions from Nrf1/2 in the transcriptional ability to regulate antioxidant response element (ARE)-driven cytoprotective genes. Collectively, an insight into Nach/CNC-bZIP proteins provides a better understanding of distinct biological functions between these factors selected during evolution from the marine bacteria to human.

**Significance:** We identified the novel ancestral subfamily (i.e. Nach) of CNC-bZIP transcription factors with highly conservativity from marine bacteria to human. Combination of bioinformatics with genomics and molecular biology demonstrated that two of the most ancestral family members classified into BATF and Jun subgroups are originated from viruses. The Jun and CNC subfamilies also share a common origin of these bZIP proteins. Further experimental evidence reveals that Nach1/2 from the marine bacteria exerts nuance functions from human Nrf1/2 in the transcriptional ability to regulate antioxidant response element (ARE)-driven genes, responsible for the host cytoprotection against inflammation and cancer. Overall, this study is of multidisciplinary interests to provide a better understanding of distinct biological functions between Nach/CNC-bZIPs selected during evolution.

## INTRODUCTION

In a changing environment, the natural evolutionary selection of all life forms, such as unicellular and multi-cellular complex organisms, ensures that only the fittest can survive and maintain a normal development and growth with a robust homoeostasis (1). These processes are thereby dependent on differential and diverse expression of optimally-selected genes. The whole life processes including the homeostatic development and growth into certain organismic bodies with distinct cellular identifications, as well as their adaptive responses to a vast variety of intracellular and environmental stresses, are tightly controlled by signaling of transcriptional networks converged on the hub activity of many different transcription factors (TFs) to regulate expression of target genes (2). Such ability of TFs is defined by binding their specific *cis*-regulatory DNA sequences e.g. antioxidant response elements (AREs) and activating protein-1 (AP-1)-binding site, in order to control the transcriptional expression of cognate genes and display relevant functional performances in many ways (3, 4).

One of the most conserved TFs is the basic-region leucine zipper (bZIP) superfamily. They are involved in the transcriptional regulation of distinct subsets of target genes by forming diverse functional homodimers and heterodimers with their partners before binding with their specific *cis*-regulatory elements (e.g. ARE or AP-1) to the gene promoter regions. The transcriptional networks formed enables the bZIP superfamily to play vital roles in cell division, proliferation, differentiation, maintenance and other processes, particularly in multicellular organisms (5). Conversely, structural and functional deficiencies in some bZIP factors can result in various diseases, including cancer, autoimmune and inflammatory diseases, and default in many other pathological processes (6–9). Furthermore, the highly conserved bZIP protein family is predominantly determined by the founding BRLZ domain that is composed of basic region (BR) and leucine zipper (LZ) within 60-80 amino acids (aa) in length (10). The basic region comprises of an approximately 16-aa consensus sequence, which is responsible for a putative nuclear localization signal (NLS) and DNA-binding to target genes. By contrast, the LZ region is composed of heptad repeats of leucine or other bulky hydrophobic residues exactly at the ‘*d’* positions, which mediates dimerization of bZIP proteins (11, 12).

With the whole genome sequences of distinct species available, ever-increasing bZIP proteins are identified as key players in defending against abiotic stresses in plants, including Arabidopsis (13), rice (14), apple (15) and maize (16). By contrast, similar transcription factors of the bZIP uperfamily existing in animals appear to have originated throughout the eukaryotic evolution process before the dawn of the Metazoa. This is supported by the fact that some of the highly conserved bZIP family proteins also emerged in the protozoa, such as choanoflagellate (*Monosiga brevicollis*) and protist (*Capsaspora owczarzaki*) (17), in addition to the presence of orthologues in the Metazoa. Immediately with accumulating analyses of the elaborate bZIP-mediated transcriptional networks within distinct eukaryotes, their evolutionary process had also been investigated in animals (17, 18), fungi (19) and plants (20). For example, the bZIP superfamily in Metazoan was thought to be evolved from a single ancestral eukaryotic gene, which had undergone multiple independent expansions and three major evolution periods. Three identifiable ancestral opisthokont bZIP proteins ATF6, ATF2-sko1 and Jun-CGN4 were found (21), but the evolutionary origin of the bZIP superfamily remains elusive.

The original orthologues of the CNC (*c*ap’*n*’*c*ollar) subfamily of bZIP proteins, including NF-E2 p45-related factors (Nrfs), has previously been considered to be the *Caenorhabditis elegans* protein Skn-1 (22), besides the founding *Drosophila melanogaster* Cnc protein (23). Fortunately, now we discover a *de facto* ancestral subfamily of CNC-bZIP proteins, that is designated “Nach” because of its high conservativity with all known Nrf/CNC proteins. Interestingly, Nach1/2 from the marine bacteria is the ancestral homologue of all other Nach/CNC proteins with the ability to regulate expression of ARE-driven genes, a distinctive role compared to Nrf1/2. Importantly, we also demonstrate that the Nach/CNC subfamily shares an early evolutionary stem with the Jun subfamily, whilst two of the most ancestral family members are originated from viruses and identified to belong to the Jun and BATF (B cell-activating transcription factor) subgroups respectively, albeit the expansion and diversification of the bZIP superfamily occurs in different vertebrates. Moreover, the membrane-bound Nach/CNC-bZIP proteins were further analyzed for their interactions with other bZIPs expressed in the human.

## RESULTS

### Species distribution and phylogenetic analysis of bZIP transcription factors

To investigate the origin of the bZIP family members, some known bZIP sequences were used as queries for BLASTP (i.e. protein blast) in the non-redundant protein sequences database and for the HMMER (i.e. Hidden Markov Model) search. The bZIP proteins that were selected span from 23 virus to mammal species (Fig. 1A). Across the 23 species, a total of 495 of the bZIP proteins were identified, after removal of both incomplete and repeated sequences from the resulting searches from BLASTP and HMMER databases. For the *Gallid herpesvirus* 2, only one bZIP protein (called MEQ) was identified, with a BRLZ domain that had a high identity of 60.82% with the BATF subfamily (Fig. S1A). Interestingly, additional two bZIP proteins were found in the marine bacteria *Endozoicomonas* (*E.*) *sp. ab112s* and *E. numazuensi*, which are designated Nach1/2, respectively, based on its high homology with the CNC-bZIP proteins (Figs. S2 to S4). In metazoans, except vertebrata, the number of bZIP proteins were approximately between 11 and 20 (Fig. 1A), for an example sea urchin up to 20 proteins. The more bZIP proteins were identified in vertebrata, e.g. African clawed frog (*Xenopus laevis*) had up to 87 bZIP proteins, 11 of which belong to the CNC-bZIP subfamily.

**Fig. 1.**
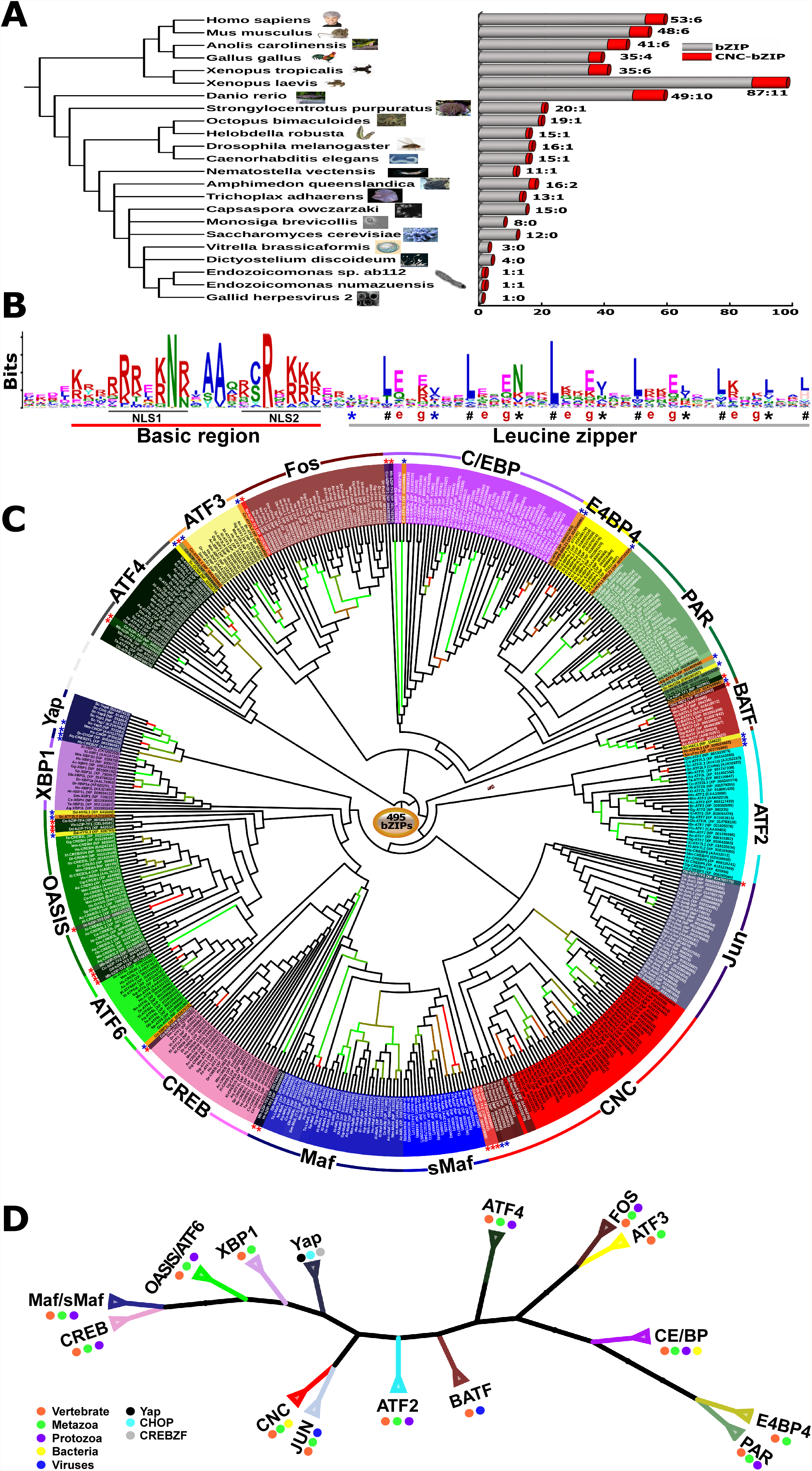
Species distribution and phylogenetic analysis of bZIP transcription factors. **(A)** *Left panel* shows distinct evolutionary status of representative species with distinct organism pictures, whilst *right graph* illustrates discrepant distribution of total bZIP *(in grey column)* and particular Nach/CNC-bZIP *(in red column)* proteins existing in each of different species. They include Gallid herpesvirus 2 (Gh2), Endozoicomonas numazuensis (En), Endozoicomonas sp.abll2 (Es), Dictyostelium discoideum (Dd), Vitrella brassicaformis (Vb), Saccharomyces cerevisiae (Sc), Monosiga brevicollis (Mb), Capsaspora owczarzaki (Co), Trichoplax adhaerens (Ta), Amphimedon queenslandica (Aq), Nematostella vectensis (Nv), Caenorhabditis elegans (Ce), Drosophila melanogaster (Dm), Helobdella robusta (Hr), Octopus bimaculoides (Ob), Strongylocentrotus purpuratus (Sp), Danio rerio (Dr), Xenopus laevis (XI), Xenopus tropicalis (Xt), Gallus gallus (Gg), Anolis carolinensis (Ac), Mus musculus (Mm), and Homo sapiens (Hs). **(B)** Shows a colour Logo image obtained from MEME analysis of both the basic-region (BR) and leucine zipper (LZ) domains within 495 of bZIP transcription factors, in which the location of a bipartite nuclear localization signal (NLS, that is composed of two parts NLS1 and NLS2 is underlined, whilst the *’a’* and *’d’* positions at the putative helixes folded by six heptad repeats are indicated by different symbols * and #, respectively. **(C)** Shows an evolutionary wheel built by the phylogenetic analysis of 495 bZIP proteins across 23 species. The more than 50% of bootstrap values were shown by green to red clades (those < 50% are not marked). **(D)** Display a simplified phylogenetic tree of all identified bZIPs transcription factors from distinct kinds of organisms. Such the phylogenetic trees were generated by using the MEGA 6.0-based bioinfomatic analysis of all BRLZ domains with the neighbor-joining (NJ) method with 1000 bootstrap replicates.

The sequences of the bZIP domains within the 495 proteins were extracted using the SMART software, and the conservatism principle is presented on the basis of the MEME analysis. The conservative domain of about 60 amino acids in length mainly includes the BR and adjacent LZ regions (Fig. 1B). The bipartite NLS-containing BR region consists of about 21 amino acids with the conserved motif -K/R-X3-(R/K)2-X-K/R-N-R/K/N-X-A/S/Y-A/V-X2-C/S-R-X-(K/R)3-(where X indicates any amino acid residues). Of note, the -C/S-R-peptide exists in an overwhelming majority of bZIP proteins, but is replaced by -A-R-in the viral MEQ and bacterial Nach1 (Figs. S1 and S2) or by -Y/F-R in the yeast activator proteins (Yaps, Fig. S5). The BR-adjacent LZ region is composed of six rounds of the conserved heptad repeats, wheeled by seven residues (denoted *a* to *g*), in which the typical residues at positions ‘*a’* and ‘*d’* are key in forming a hydrophobic interface essential for the dimerization of related bZIP proteins. Within the LZ regions, almost all the ‘*d’* positions are highly conserved and occupied by leucines (L) or other hydrophobic residues. The third ‘*a’* position asterisked is also highly conserved by asparagine (N), whilst glutamic acids (E) at the first to fourth ‘*g’* positions are relatively conserved (Fig. 1B).

To further clarify the evolutionary phylogenetic relationships of the 495 bZIP proteins, based on the sequence conservativity of their BRLZ domains, we constructed the neighbor-joining phylogenetic tree with 17 distinct clades subdivided from the stem as wheeled (Fig. 1C). Notably, the unicellular yeast Yaps, together with other five multi-cellular Metazoan bZIP proteins including Hs-CREBZF [cyclic AMP response element (CRE)-binding protein (CREBP)-Zhangfei], Aq-CREBZFL, Hs-CHOP [CCAAT/enhancer-binding protein (C/EBP)-homologous protein], Mm-CHOP and Dr-CHOP were gathered into a branch Yap subfamily of bZIP transcription factors, by employing the well-supported bootstrap values. The remaining 484 BRLZ sequences are clustered into additional 16 branches, each of which has a potential individual difference from others, within their mutual relatively independent and interrelate connective evolutionary trajectories (Fig. 1C). According to the clockwise direction, the gap in the phylogenetic tree wheel is referred to as a starting point, followed by distinct clusters of ATF4, ATF3, Fos, C/EBP, E4BP4 (E4 promotor-binding protein 4), PAR (proline- and acid-rich bZIP), BATF, ATF2, Jun, CNC, Maf (musculoaponeurotic fibrosarcoma oncogene homolog), sMaf (small Maf), CREB, ATF6, OASIS (old astrocyte specifically-induced substance, also called CREBP3-like protein 1), and XBP1 (X-box binding protein 1) subfamilies (of which all their BRLZ sequences were also aligned, as shown in supplemental Figs. S3 and S4, S6 to S13). Of note, Mb-ATF4L is the most primitive homologue amongst the ATF4 subfamily comprising 27 member proteins (Fig. 1C and Fig. S9B). Both the ATF3 and Fos subfamilies appear to serve as two homogeneous descents conjunctly from a big predecessor branch (Fig. 1C), whilst another large progenitor branch is jointly gathered together with the E4BP4/NFIL3 and PAR subfamilies that are clustered with the big C/EBP subfamily clade. Additional 3 antecedent branches are gathered phylogenetically with each of two descent subfamilies Jun and CNC, the Maf (that is combined with sMaf) and CREB, OASIS and ATF6 respectively, besides others subgroups clustered independently.

To simplify the above-described phylogenetic tree, the trunk with distinct evolutionary branches representing different subfamilies was collapsed (Fig. 1D), particularly with distinct species lineages marked for the existence of different bZIP subfamilies. More interestingly, both the Jun and BATF subfamilies are inferable to be originated from the earliest primogenitor existing in the viruses (Fig. S1A and B), whereas either the C/EBP (Fig. S1C) or CNC (Figs. S2 to S4) subfamilies appear to be stemmed from the ancestral homologues emerged in the bacteria. Furthermore, both the XBP1 and E4BP4 subfamilies seem to have early appeared in the metazoans (Fig. 1D), whilst all other nine subfamilies Maf/sMaf, CREB, OASIS/ATF6, ATF2, ATF4, FOS and PAR have surprisingly shared with those derivatives from the putative primogenitor-originated protozoans selected before the dawn of the metazoans (also see Figs. S6 to S13). In addition, there exists a commonly-sharing predecessor of yeast Yap proteins, metazoan CREBZF and CHOP subgroups.

### A novel evolutionary branch of the CNC-bZIP subfamily from ancestral Nach proteins

According to the current literature, we have found that the CNC subfamily of bZIP transcription factors is composed of NF-E2 p45, Nrf1, Nrf2, Nrf3 and their transcriptional repressors Bach1 and Bach2 in vertebrates, in addition to Cnc and Skn-1 proteins found in *Drosophila* and *Nematodes*, respectively. To investigate the evolutionary origin of these CNC-bZIP proteins, a new line of the inquiry sequences of the human orthologous molecules were employed for the BLASTP in the non-redundant protein sequences database and for the HMMER search. As shown in Fig. 2A, it was discovered that, apart from Skn-1 and Cnc in *Caenorhabditis elegans* and *Drosophila melanogaster*, the most original homologies with the CNC-bZIP proteins are objectively present in the marine bacteria *E. numazuensis* (Nach2) and *E. sp.ab112* (Nach1). More excitingly, other proteins homologues with CNC-bZIP were further searched in the primitive multicellular organism *Amphimedon queeslandica* (Nach4/5) and also discovered in the following invertebrates *Trichoplax adhaerens* (Nach3), *Nematostella vectensis* (Nach6), *Octopus bimaculoides* (Nach7) and *Strongylocentrotus purpuratus* (Nach8) (*Left panel*). Hence, based on these ancestral bZIP homologues at their evolutionary status selected from the origin to the advance in the diversity of such bZIP-distributed species, they are herein classified into a novel subgroup that is nominated as Nach (*N*rf *a*nd *C*NC *h*omology) (i.e. Nach1 to Nach8, Fig. 2A, *right panel*). In addition, it should be noted that amongst vertebrates, *Danio rerio* and *Xenopus laevis* have given rise to 10 and 11 of CNC-bZIP homologues, respectively, but only 4 of CNC-bZIP proteins exist in *Gallus gallus* with a constructive loss of NF-E2 p45 and Nrf3 from within the genome.

**Fig. 2.**
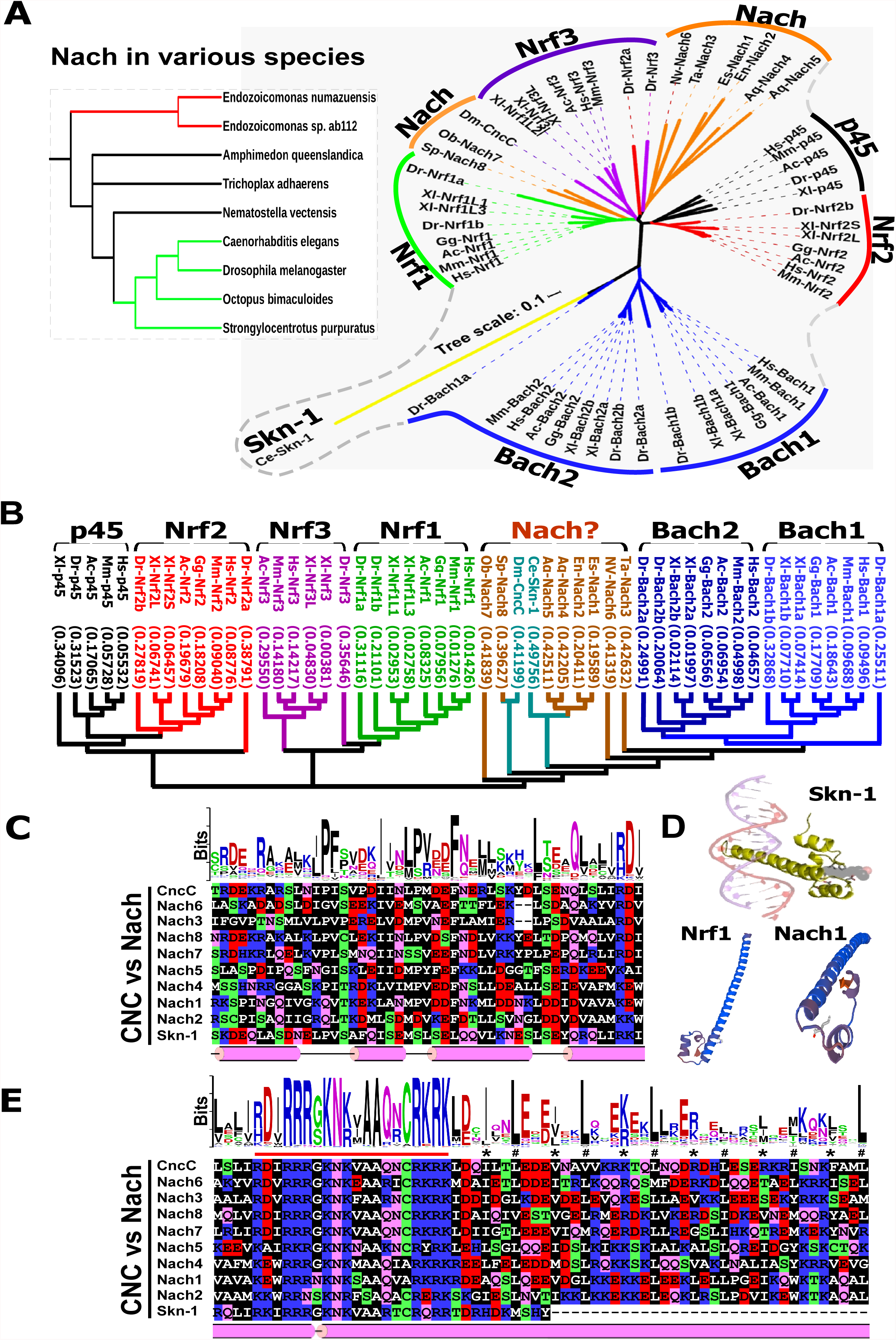
Phylogenetic analysis and sequence structure of CNC-bZIPs in various taxas. **(A)** *Left panel* shows the putative evolution of a novel clade called the Nach subgroup form distinct organisms, which are located in the *right* smaller fan-like phylogenetic tree that was generated by using the MEGA 6.0-based analysis of the full-length Nach/CNC-bZIP proteins across 17 different species with the neighbor-joining method (with 1000 bootstrap replicates). (**B)** Another phylogenetic tree was generated by using the T-coffee analysis of the above full-length Nach/CNC_bZIP proteins with the default parameters. **(C** *to* **E)** Multiple sequence alignments of both CNC **(C)** and BRLZ (£) domains with distinct characteristics were analyzed by using different tools DNAMAN8.0, PSIPRED, MEME and Web-logo with distinct default parameters. Two similar secondary (C, *E on the bottoms)* and tertiary **(D)** structures of the CNC-BRLZ domains within Nrf1 and Nach1 were modeled by using the SWISS-MODEL tool, on the temple of the known homological domain structure of Skn-1.

By bioinformatic analysis of CNC-bZIP full-length amino acid sequences, 60 of these proteins that have been identified so far were further sub-clustered in order to construct a smaller neighbor-joining phylogenetic tree (Fig. 2A). Distinct blades of NF-E2 p45, Nrf1, Nrf2, Nrf3, Bach1 and Bach2, together with two Nach blades separated by Nrf3, are all displayed in a fan-like phylogenetic tree that is protruded with a seemingly longer evolutionary branch of Skn-1 in nematodes (*right panel*). This is due to a genomic loss of the zipper domain within Skn-1, albeit it retains both its basic region and CNC domain. Moreover, an additional type of similar phylogenetic tree was also constructed by the T-Coffee software, such that a single typical clade of Nach proteins was shown clearly (Fig. 2B).

The multiple sequence alignment of the above-identified CNC-bZIP proteins revealed one of the most conserved motifs, -ϕ^10^-φ-I/L-P/Q^13^-F/ϕ-X2-φ2-I/L-ϕ/T^20^-φ-L/M-P/S^23^-V/R^24^-φ-D/E-F-N/Q-X-ϕ2-x4-L/F-X3-Q/φ-ϕ-X-ϕ-ϕ^44^- (in which ϕ and φ represent any hydrophobic and hydrophilic residues respectively, besides X denoting any aa) within their CNC domains (Fig. 2C and Fig. S3). Notably, a remarkable difference between the Bach1 and Bach2 subgroups appears to be made by the latter 20^th^ position-specific threonine (T, in Bach2) or the former 23^rd^ position-specific serine (S,in Bach1), in addition to their 24^th^ position-specific arginines in both subgroups, which are strikingly distinctive from any hydrophobic residues occupying at these same corresponding positions in all other CNC-bZIP subgroups. Overall, the motif is highly conserved by its sequence identity with equivalents existing amongst those advanced eukaryotes from octopus to humans, but in some lower lineages, it is a relatively less conservative across all their CNC domains. For example, glutamine (Q) is occupied specifically at the 13^th^ position of the bacterial Nach1 or Nach2 proteins, but is replaced by proline (P) within almost all other CNC-bZIP proteins (Fig. S3). Moreover, the secondary structure of this domain is folded into three α-helixes and the N-terminal part of the fourth α-helix, all of which were separated by linear coils (Fig. 2C and D).

Similarly, their BRLZ domains also contain the typical basic region and leucine zipper within all CNC-bZP proteins (Fig. 2E). There are highly conserved 21-aa residues -B-D/E-ϕ-R3-G/S-K-N-K/R-ϕ-A2-Q/R-N/K-C-R-K-R-K-ϕ- (in which B indicates a basic residue) in the basic region. By contrast, the less conserved leucine zipper is composed of six heptad repeats of leucines at the ‘*d’* positions in the α-helical coiled coil as wheeled (Fig. 2D and E), in which a high consistency of the first to the third repeated leucine residues (and also the last one) is maintained in all CNC-bZIP proteins except Skn-1 (lacking this LZ domain), but the remaining fourth and fifth repeats are less conservative (also see Fig. S4). In addition, the homology modeling of the CNC and adjacent BRLZ domains in Skn-1, Nrf1 and Nach1 by the SWISS-MODEL tool predicted that the latter two proteins have a similar three-dimensional structure to the known template of Skn-1 and other bZIPs (Fig. 2D).

### Distinct subgroups of the membrane-bound bZIP transcription factors

Bioinformatic analysis using the TMpred and TMHMM tools of potential transmembrane (TM) domains within the above-identified bZIP proteins across 23 species, has predicted that the putative TM domains are present in the bZIP factors, OASIS, ATF6, XBP1, CNC and other ambiguous proteins (Fig. 3A), along with the known membrane-bound SREBPs (Sterol-regulatory element binding proteins), which acts as a basic helix-loop-helix zipper (bHLH-ZIP) transcription factors, and also contains two TM domains denoted SREBP-TM1 and -TM2 from its N-terminal to its C-terminal ends, respectively. These possible TM sequences were manually used for further analyses of their hydrophobicity and evolutionary conservativity. The resulting phylogenetic tree with several branches is extended to distinct four major clades, of which the same categories gathered together were collapsed before being separately displayed in details somewhere else (Fig. 3A, *cf. left with right panels*). Notably, two small subgroups of XBP1 and CNC-NHB1 [in which NHB1, i.e. the N-terminal homology box 1, was identified to enable its related CNC-bZIP proteins to be integrally anchored within the endoplasmic reticulum (ER) (24, 25)] were gathered conjunctly to a branch. Whilst the SREBP-TM1 and the majority of OASIS and ATF6 were clustered into a large class, additional classes are gathered together with the remaining few of OASIS and ATF6, plus the SREBP-TM2 subgroup (Fig. 3A). Interestingly, the last branch of mouse Nrf1D (i.e. a variant of Nrf1) (26) embraces with other bZIP proteins from different subfamilies (Fig. 3A, *left bottom*).

**Fig. 3.**
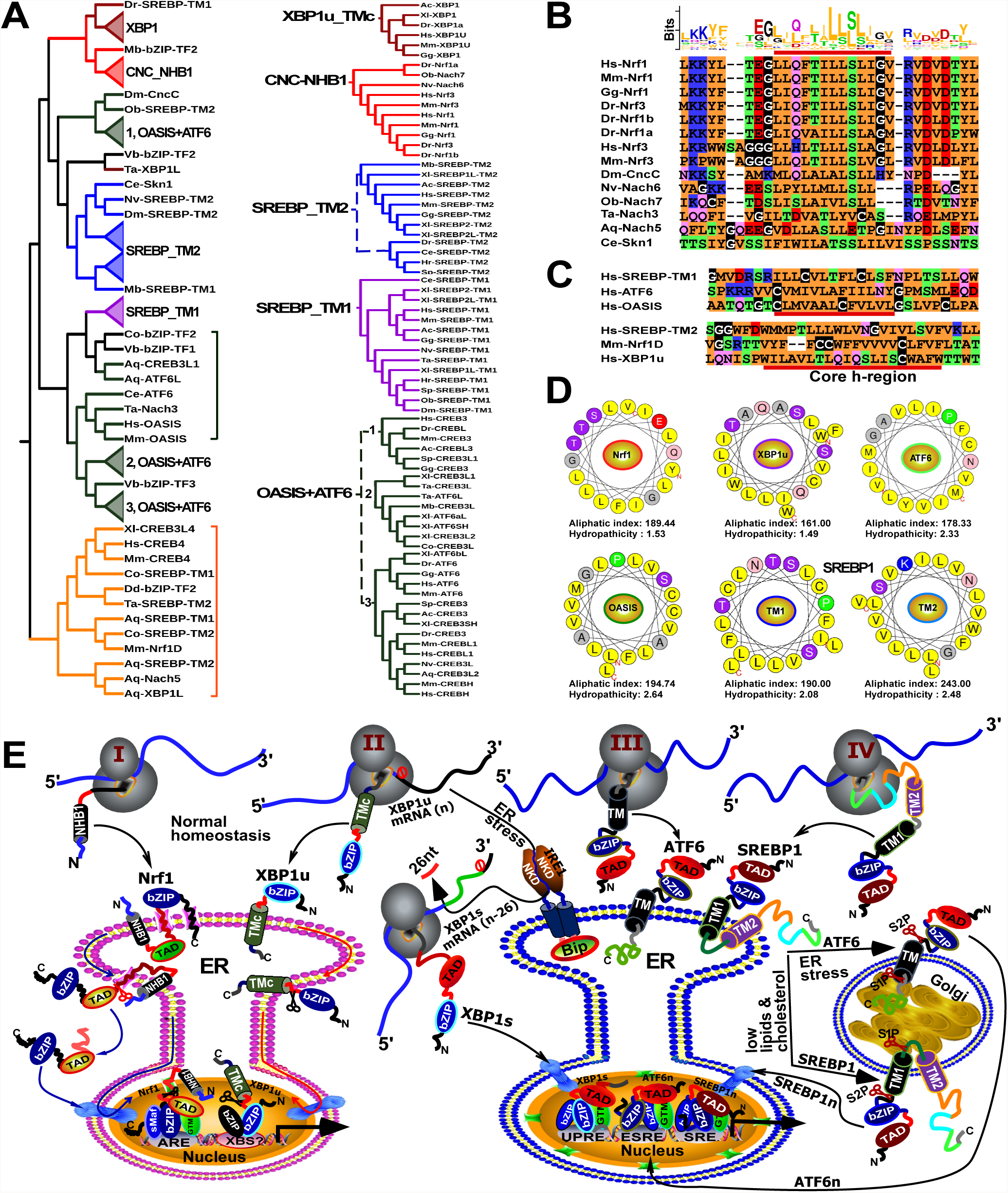
Classification of the TM-containing transcription factors. **(A)** All the putative TM-containing domains within bZIP and SREBP proteins were subjected to construction of the neighbor-joining phylogenetic tree by using the MEGA 6.0 analysis with 1000 bootstrap replicates. **(B)** A multiple sequence alignment, with a color Logo of those NHB1-associated TM regions within related Nach/CNC-bZIP proteins, was carried out by using the DNAMAN8.0 and MEME tools. **(C)** Shows two similar sequence alignments of additional TM domains of ATF6, OASIS, Nrf1D and XBP1u with SREBP1-TM1and -TM2, of which the hydrophobic h-region cores are underlined. **(D)** Six wheels of a-helixes are folded by the NHB1 – associated TM1 of Nrf1, the C-terminal TM region of XBP1u, the central TM domains of ATF6 and OASIS, as well as SREBPl-TM1 and -TM2, respectively. Both aliphatic index and the hydropathicity were also calculated. **(E)** Four distinct membrane-topobiology models are proposed to give a clear explanation of the TM-containing transcription factors with discrepant catalogues of dynamic topological folding within and around the ER and/or Golgi apparatus, before being retrotranslocated out of membranes in order to be released and transferred into the nucleus, prior to transactivating different sets of target genes, under normal homeostatic or the ER-derived stress conditions.

According to the current topological knowledge (27, 28), it is plausible that the *bona fide* TM domains are dictated by their constitutive core hydrophobic (h)-region spanning across membranes, whilst distinct orientations of these segments within membranes are predominantly determined by charge differences between its n-region and c-region flanking the core h-region. Therefore, further multiple sequence alignment of the putative TM domains in the CNC-NHB1 subgroup (Fig. 3B), the C-terminal TM domain (distinctive from its N-terminal NHB1) of mouse Nrf1D, human SREBP-TM1, SREBP-TM2, and others from human ATF6, OASIS and XBP1u (Fig. 3C) revealed that they are indeed composed of the major hydrophobic residues through their core h-regions. In particular, almost identical sequences of the core h-regions were presented in the vertebrate CNC-NHB1 subgroup (Fig. 3B). By contrast with NHB1, a relative less conservative TM domain was also observed in each of the four novel Nach3, 5, 6 and 7 proteins. Furthermore, although the C-terminal TM regions from both mouse Nrf1D and human XBP1u share a certain conservativity with human SREBP-TM2 domain, they also share a relative poor consistency in their sequences as aligned (in Fig. 3C, *lower panel*), but an additional highly consistency is provided by SREBP-TM1 and homologies from the majority of OASIS and ATF6 subgroups (*upper panel*).

Subsequently, in order to give a clear explanation of topological folding of the putative membrane-proteins, such typical TM α-helixes of Nrf1 (as a major representative of the CNC-NHB1 subgroup), XBP1u, ATF6 and OASIS, as well as SREBP-TM1 and -TM2, were herein wheeled by the HeliQuest tool, with their aliphatic indexes and hydropathicity estimated (Fig. 3D). These six α-helixes are indeed endowed with highly estimated values, of which SREBP-TM2 possesses the highest aliphatic index up to 243, whilst the highest hydropathicity of OASIS’ TM region is up to 2.64. Further, the above-predicted membrane-bound (bZIP and bHLH-ZIP) transcription factors were summarily classified into four different categories, which are Nrf1, XBP1u, ATF6 and SREBP1, to provide a better understanding of distinct topovectorial folding processes of the putative TM-containing proteins integrated within and around membranes (Fig. 3E). Firstly, the N-terminal SPase-uncleavable NHB1 signal sequence of Nrf1 enables it to be integrally anchored within the ER membranes and determines its topological folding of adjacent domains and their partitioning into the luminal or cytoplasmic sides of membranes (Fig. 3E, *Model 1*) (29). Subsequently, dynamic repositioning of the luminal-resident transactivation domain (TAD) of this CNC-bZIP factor is driven by p97-fueled retrotranslocation pathway into the cytoplasmic side of membranes. Its deglycoprotein is therein allowed for further proteolytical processing by cytosolic proteases to yield a cleaved mature factor in order to be released from membranes and translocated into the nucleus before the formation of a functional heterodimer with its partner sMaf or other bZIP proteins, which ensures its different transcriptional regulation of ARE-driven genes (26). Secondly, the unspliced XBP1u mRNA and its protein are targeted to the ER membrane (30–32). Under the normal conditions, the prototypic XBP1u protein is also anchored within membranes through its C-terminal TMc region (Fig. 3E, *Model 2*), in a topology similar to that of the C-terminal Nrf1D, before eliciting its unique function as a transcriptional repressor. Upon the exposure to ER stress, the splicing of XBP1u mRNA by IRE1 to remove 26 nucleotides results in an open reading frame-shifting variant XBP1s, which lacks an available TM-targeting peptide so as to directly translocate the nucleus and regulate target genes involved in the ER-to-nuclear unfolded protein response (UPR) (*right panel*). Thirdly, ATF6 is folded to adapt its initial membrane-topology within and around the ER, but stimulation by ER stress enables it to be transported to the Golgi apparatus (Fig. 3E, *Model 3*), in which this protein is allowed for the progressive two-step processing of it by Site-1 and Site-2 proteases (i.e. S1P and S2P) (33, 34) to give rise to a cleaved active factor ATF6n before transactivating its downstream genes driven by ESRE (ER stress response element) and/or UPRE (UPR element) within their promoter regions. Finally, although SREBP1 encompasses two TM domains with distinct local topologies integrated within and around the ER, only its TM1 is folded in a similar orientation to that of ATF6 such that this bHLH-ZIP protein, only when its target genes are required for cholesterol and other lipid synthesis, undergoes a similar transfer from the ER through the Golgi to the nucleus (Fig. 3E, *Model 4*). This process is also attributed to regulated intramembrane proteolysis of the protein by SIP and SIP2 successively in the Golgi apparatus to generate a cleaved activator SREBP1n (34, 35).

### Expressive differentiations of human bZIP factors within their interaction networks

To gain an insight into the evolutionary diversity of human bZIP subfamilies during the process of phylogenesis, their neighbor-joining tree was constructed on the base of all their full-length sequences (Fig. 4A). The phylogenetic tree displays six major branches including 17 minor subgroups, except that CREBZF is clustered separately. Amongst them, there are high bootstrap values in the nodes of between CNC and Jun (0.61) or between ATF3 and Fos (0.98), besides sMaf and Maf belonging to the same large category, whilst other three subgroups of ATF6, OASIS and CREB are also homologous. Next, in order to determine potential biological functional differentiation of these human bZIP genes, their mutual interaction networks were established on the ground of experimentally validated evidence, by using the STRING program of 51 bZIP-interactive proteins (with a score > 0.7 of the moderate confidence) (Fig. 4B and C). At the center of the interaction network, the CNC-bZIP subfamily proteins have indeed frequent networking with sMaf, and additional complex links also exist between other bZIP proteins. These two distinct expression profiles of human bZIPs were obtained from bioinformatic analysis of the total RNA sequencing datasets. When *Nrf1α^−/−^* was knocked out by Talens-mediated editing of its genome in the liver cancer HepG2 cells (36), at least 16 bZIP factors, such as *Bach2, MafK, MafF, Jun, FosB, Fra1, ATF3, ATF4, NRL, HLF, TEF, CREB5, CEBPE, E4BP4*, *BATF3* and *CREM* were up-regulated significantly (with > +1 of the Log2-based RPKM value calculated) (Fig. 4B and D), whereas other 7 bZIP factors *NF-E2 P45, MafA, JunD, DBP, CEBPA, CEBPD*, and *BATF2* were down-regulated significantly (with < −1 of the Log2-based RPKM value), in addition to *BATF, MAF* and *CREBH* possibly at much lower levels such that they were not detected by RNA-sequencing. Conversely, the stable tetracycline-inducible expression of Nrf1α in HEK293C^Nrf1α^ cells (Fig. 4C and D) caused significant increases in the abundances of *Nrf2, ATF2, CREB1, DBP* and *CREB5*, while additional expression of *Fos, CEBPE* and *BATF2* were significantly weakened, but both *BATF* and *CREBH* were also not detected, implying distinct cell-specific expression of certain bZIP genes possibly in different biological processes.

**Fig. 4.**
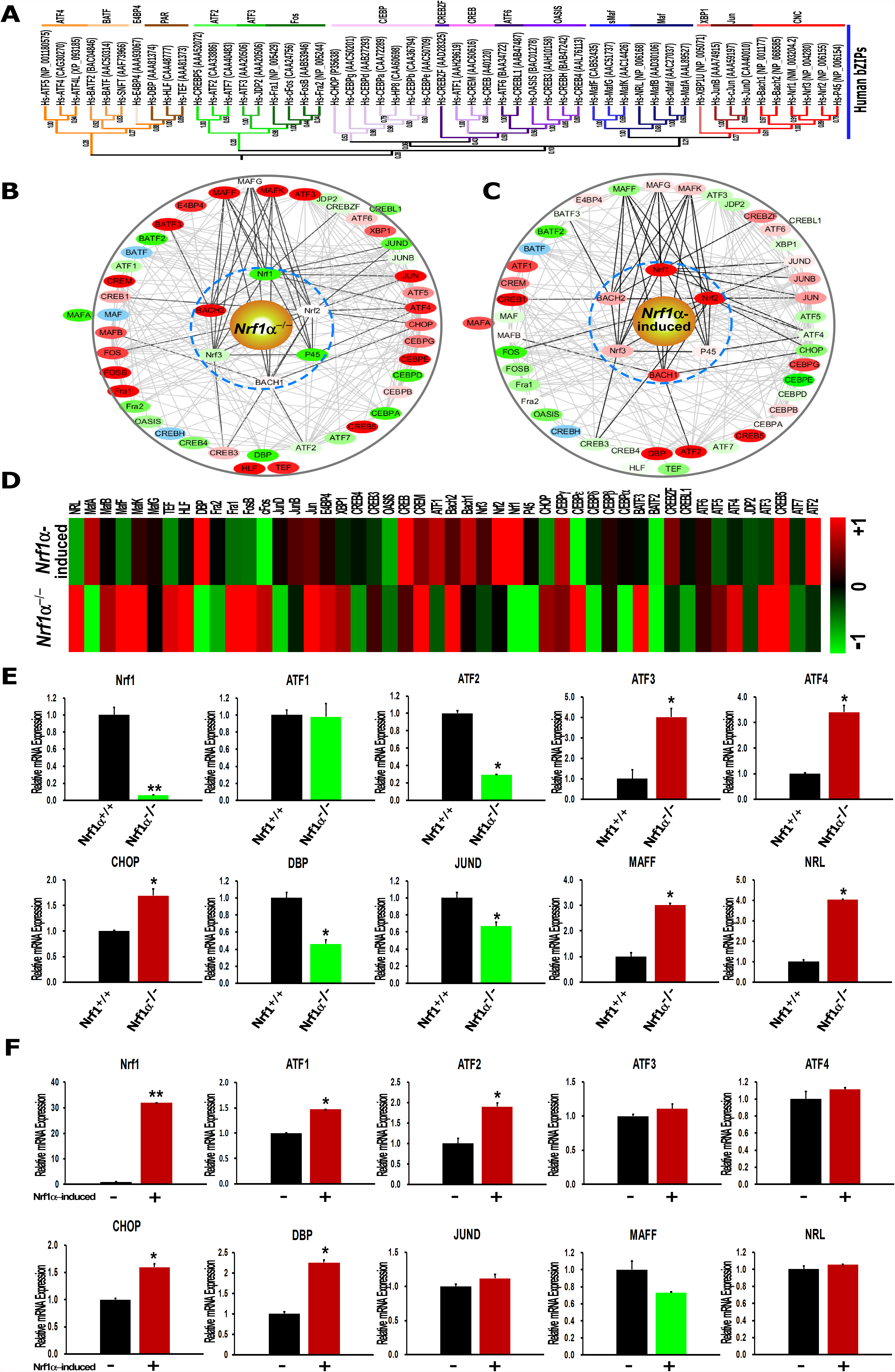
Classification of human bZIP proteins within their interaction networks converged on a hub of Nrf1α. **(A)** The phylogenetic tree of 53 bZIP proteins in humans was constructed by the same method as described in Figure 2. **(B, C)** Distinct or opposing changes in some nodes within the two interaction networks, which are composed of all human bZIP proteins and also converged on a hub of Nrf1α, were determined following knockout of *Nrf1α*^−/−^ **(B)** or induction of this protein expression by tetracycline in HEK293C^Nrf1α^ **(C).** Significant up-regulation (Log2-based RPKM value >1) of the indicated genes were red-labeled, whereas down-regulation (Log2-based RPKM value <–1) of other indicated genes were green-labeled. Such a green-to-red gradient of those coding genes demonstrates from being down-to up-regulated. Additional genes without any detectable signals by sequencing were also blue-labeled. **(D)** A heat map was made by the Log2-based RPKM values, which represents differential expression profiles of human bZIP proteins in *Nrf1α*^−/−^ or HEK293C^Nrf1α^ (when compared to *Nrf1^+/+^* HepG2 or un-stimulated HEK293C cells, respectively). Different changes in the expression of some genes were shown to distinct degrees of colourings scaled. **(E, F)** Relative expression levels of selected bZIP genes were also validated by qRT-PCR analyses of *Nrf1α^−/−^ Nrf1^+/+^* (*E*), or the stable tetracycline-inducible HEK293C^Nrf1α^ *vs* unstimulated cells (*F*). Subsequently, significant decreases (*p<0.05, **p<0.01) or increases ($, p<0.05; $$, p<0.01) in the expression of some genes were determined.

The expression profiles of some key genes selected from the above RNA-seq datasets were further validated by quantitative real-time PCR (qRT-PCR) analysis of the following 10 bZIP factors. In *Nrf1α^−/−^* knockout cells (Fig. 4E), *ATF3, ATF4, CHOP, MafF* and *NRL* were significantly up-regulated at their mRNA levels (*P*<0.05), whereas *Nrf1, ATF2, DBP* and *JunD* were markedly down-regulated (*P*<0.05), but ATF1 was unaltered. By contrast, induction of Nrf1α by tetracycline in HEK293C^Nrf1α^ cells resulted in a significant up-regulation (*P*<0.05) of *Nrf1, ATF1, ATF2, DBP and CHOP* (Fig. 4F), but only caused a marked down-regulation of *MafF* alone (*P*<0.05), in addition to no obvious changes in the expression of *ATF3, ATF4, JunD* and *NRL*.

### Nach1 shares conserved domains with other CNC-bZIP factors at differently regulating target genes

For an in-depth insight into functional domains of Nach1 and Nach2 required for the regulation of ARE-driven genes (as evidenced by its orthologues of the CNC-bZIP proteins Nrf1, Nrf2, NF-E2 p45) (26), their conserved structural domains were presented schematically (Fig. 5A). The schematic shows that the bacterial Nach1 shares similar structural domains closer to those of human NF-E2 p45 than Nrf1 and Nrf2, demonstrating that p45 is preserved as a possible reminiscent of its evolutionary process from the progenitor factor Nach1. By sharp contrast, the ancestral Nach1 lacks the Neh5L domain (Fig. 5A and Fig. S2B), which is essential for transactivation of ARE-driven genes by all known CNC-bZIP activators (29, 37, 38), and is hence postulated to act as just a transcriptional repressor as Bach1. The putative negative regulatory function of Nach1 was also further inferable to be monitored through its potential degron DSGxSL (i.e. canonical) and another similar motif DSGxxL (i.e. non-canonical), both with distinct locations from equivalents of NF-E2 p45, Nrf1 and Nrf2 (Fig. 5A and Fig. S2G), whilst Nach2 only retains the canonical DSGxSL degron in a similar location to that of Nach1.

**Fig. 5.**
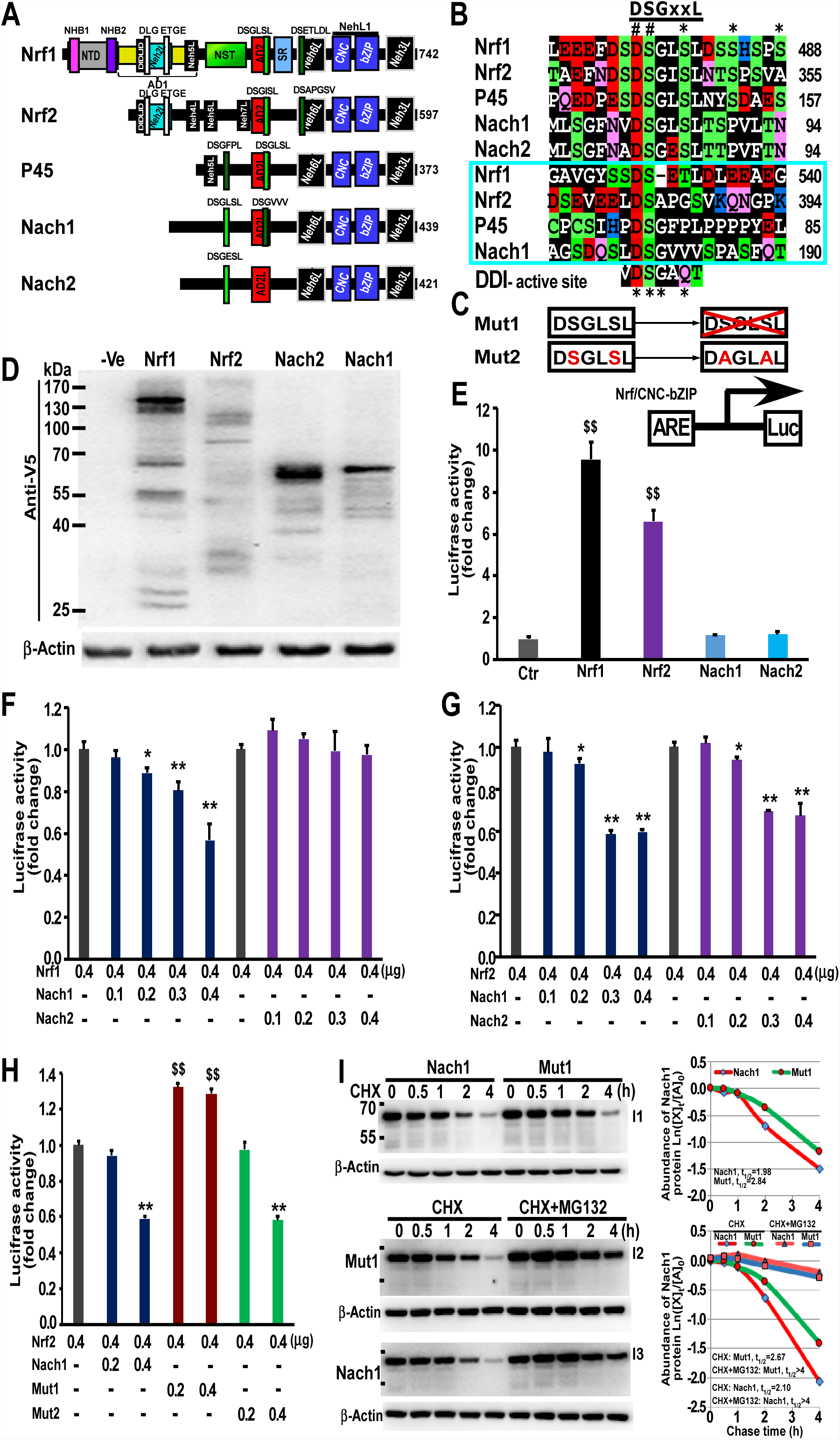
Distinctions in ARE-driven gene regulation by Nrf1 and Nrf2 from Nach1 and Nach2. **(A)** Schematic represents structural domains of Nrf1, Nrf2, NF-E2 p45, Nach1 and Nach2, in which the locations of a canonical DSGxSL degron and another non-canonical DSGxxL motif are indicated. **(B)** Shows a multiple alignment of the DSGxSL and DSGxxL [*boxed*)-adjoining sequence within Nrf1, Nrf2, NF-E2 p45, Nach1 and Nach2. The two degron motifs are highly conserved with the enzymatic active sites (DSGxQx) of the DDI aspartic proteases and hence also denoted as a putative suicidon. **(C)** Diagrammatic represents the DSGLSL motif and its mutants of within Nach1. **(D)** Western blotting of HepG2 cells that had been transfected with an expression construct for C-terminally V5-tagged Nrf1, Nrf2, Nach1 and Nach2, whilst β-Actin served as an internal control to verify amounts of proteins that were loaded in each well occasion. **(E)** ARE-driven Luciferase reporter gene activity was measured from HepG2 cells that had been transfected for 24 h with an expression construct (0.4 μ**g of cDNA)** for Nrf1, Nrf2, Nach1 and Nach2, together with *GSTA2-6xARE-Luc* plasmid (0.2 μ**g)** and pRL-TK (0.1 μ**g**, as an internal control), and allowed for a 24-h recovery from transfection before being disrupted in the lysis buffer. The data were calculated as a fold change (mean ± SD) of transactivation by indicated Nach/CNC-bZIP factors. Significant increases ($, p<0.05; $$, p<0.01) in the transactivation activity of reporter gene are determined relatively to the control value. **(F, G)** Additional two measures of ARE-driven luciferase reporter activity was carried out in HepG2 cells that had been transfected with an expression construct (0.4 μ**g of cDNA)** for Nrf1 (*F*) or Nrf2 (G) in a combination with distinct concentrations (from 0.1 to 0.4 μ**g)** of either Nach1 or Nach2 expression plasmids, together with *GSTA2-6xARE-Luc* plasmid (0.2 μ**g)** and pRL-TK (0.1 μ**g)** as described above Significant decreases (*, p<0.05; **, p<0.01) in the reporter activity are indicated relatively to control. (H) Similar luciferase reporter activity was also determined in HepG2 cells that had been transfected with 0.4 μ**g** of an expression construct for Nrf2 alone or plus another expression construct (0.2 to 0.4 μ**g)** for Nach1 or its mutants, along with *GSTA2-6xARE-Luc* plasmid (0.2 μ**g)** and pRL-TK (0.1 **pg).** The resulting data were calculated as described above (I) HepG2 cells that had transfacted with an expression construct for Nach1 or Mut1 were treated with CHX (50 μg/ml) alone or plus MG132 (10 μmol/L) for 30 min to 4 h before being disrupted. The relative expression levels of Nach1 or Mut1 in the total lysates were determined by Western blotting with ant -V5 antibody (*left panels*). The intensity of indicated proteins with distinct half-lives was quantified and shown graphically [*right panels*).

To determine the biological function of Nach1 and Nach2, HepG2 cells were transfected with each of expression constructs for Nach or CNC-bZIP protein that is tagged C-terminally by the V5 ectope. Western blotting showed that either Nach1 or Nach2 was expressed as a major protein of 65-kDa estimated on 10% PAGE gels (Fig. 5D), which was also accompanied by a ladder comprising of several degraded polypeptides between 65-kDa and 30-kDa, indicating that both proteins are unstable and rapidly destructed. A luciferase assay revealed that over-expression of Nach1 or Nach2 only caused a marginal activity of ARE-driven reporter gene, increased by 15.33% and 21.53% of, despite no extremely significant differences from, the background levels (Fig. 5E), when compared with ectopically expressed Nrf1 and Nrf2 leading to significant increases in the transactivation of this reporter gene by 9.56- or 6.60-fold, respectively. However, co-transfection experiments showed that Nach1, but not Nach2, caused a marked decrease in the Nrf1-mediated transactivation activity by 11.19% to 43.33% in a dose-dependent manner (Fig. 5E). Similar dose-dependent inhibitory effects of Nach1 were also exerted on the Nrf2-mediated transactivation activity so as to be decreased by 7.77% to 40.47% (Fig. 5F), whilst another reduction in this activity by 6.14% to 32.56% also resulted from co-expression of Nach2 with Nrf2.

Next, to identify a role of the putative DSGxSL degron within Nach1, its mutant 1 (i.e. Mut1, which was yielded by mutagenesis to delete the entire DSGLSL motif from Nach1) and Mut2 (in which the DSGLSL motif was mutated to DAGLAL) were subjected to co-expression with Nrf2 in ARE-driven reporter assays (Fig. 5C and H). As anticipated, Mut1 rather than Mut2 resulted in a striking de-repression of Nrf2 by Nach1, such that Nrf2-mediated reporter gene activity was significantly restored to 1.41~2.19-fold transactivation (Fig. 5H). Further pulse-chase experiments of cells that had treated with cycloheximide (CHX, that inhibits biosynthesis of nascent proteins) alone or in combination with the proteasomal inhibitor MG132 revealed that Mut1 caused an obvious increase in the mutant protein stability with its half-life determined to be 2.84 h (=170 min) (Fig. 5I, *I1 and upper graph*), when compared with wild-type Nach1 protein with a shorter half-life defined to be 1.98 h (=119 min) following CHX treatment. However, the turnover of Nach1 and Mut1 proteins was still prolonged by MG132, with similar half-lives estimated to be over 4 h (*I2, I3 and lower graph, and also see the whole gel images in Fig. S14*).

## DISCUSSION

During the evolutionary process along with natural section, the diversity of living organisms increases their biological complexity of distinct species to survive in a changing environment. These organisms have been evolutionarily selected for distinct graded requirements of more transcription factors in order to monitor differential expression of their different sets of target genes that are involved in the normal homeostatic development and growth, and also are responsible for diverse patho-physiological adaptation and cytoprotection against a vast variety of stresses (18). In fact, it is found here that disparate species lineages are represented by divergent distribution of bZIP transcription factors in 15 representative metazoans, 5 typical protozoans, 2 bacteria and 1 virus. The number of bZIP proteins increases with increasing morphological and behavioral complexities in distinct vertebrates (e.g., 49 and 53 of bZIP proteins have been identified in zebrafish to human respectively), but relatively less are present in protozoa including choanoflagellates and fungi (Fig. 1A). This fact has demonstrated that the evolution of distinct species is positively correlated with expansion and diversification of the bZIP superfamily. Throughout the metazoan evolutionary process, all their bZIP proteins are conserved at certain extents, albeit many of their orthologues and paralogues have been endowed with strikingly different interactive specificities as described (39). This notion is also supported by further bioinformatic analyses of the consensus BRLZ domains from distinct bZIP subfamilies (Figs. S4 to S13), but their conservativity was not elucidated in these six main eukaryotic lineages including Holozoa, Fungi, Amoebozoa, Plantae, Heterokonta and Excavata (21).

Within all distinct subfamily of bZIP proteins, such a conservative domain is composed of both BR and LZ regions (Fig. 1 and Figs. S4 to S13). The dimeric specificity and stability of bZIP proteins are dictated principally by leucine and other hydrophobic residues occupied at these two ‘*d’* and ‘*a’* positions of heptad repeats wheeled, respectively (40, 41). When charged residues are placed at the ‘*a’* position, they are conferred to drive heterodimerization of bZIPs in Arabidopsis thaliana, whereas asparagine residue at the ‘*a’* position develops a tendency to form a homodimer (42). Here we observed that besides such leucine residues at the last ‘*d’* position preserved in these four subfamilies of CNC, Maf, ATF6 and OASIS, the histidine residues at this position are also conserved in another five subfamilies of Fos, Jun, ATF2, ATF3 and BATF, whilst the poor conservativity occurs in other subfamilies, implying that it is responsible for distinct specificity of their dimmerization. Additional potential differences amongst distinct bZIP subfamilies are also postulated to determine their dimeric stability. This is evidenced by the finding that the asparagine residue at the third ‘*a’* position is highly conserved in most bZIP proteins because it can elicit a limitation of LZ dimerization (43, 44). It is necessary to gradually trace divergence of such a large bZIP superfamily tree from the origin with distinct graded roots, through further evolutionary analysis of all these clusters of bZIP proteins into 17 clades (Fig. 1), of which 16 typical subfamilies were subjected to the coiled-coil arrays in humans (41), except for an extra classification in yeast Yaps. Notably, it is discovered that i) a common ancestor of between E4BP4, PAR, Fos and ATF3 subfamilies was traced back to protozoa (i.e. *Capsaspora owczarzaki*); ii) the ancestral proteins Nach1/2 (with a high homology closer to the CNC subfamily) and another homologues of C/EBP are present in the two different lineages of marine bacteria *Endozoicomonas numazuensis (or sp.ab112*) and *Endozoicomonas arenosclerae*, respectively. Of note, an amino acid sequence consistency of 72.22% between BRLZ domains of human C/EBP and its ancestral homologous protein (with a GenBank accession No. WP_062270874) is determined (Fig. S1C); iii) an original homologous protein of Jun has existed in *Cyprinid herpesvirus 1* (with a GenBank No. YP_007003813), with an 87.50% BRLZ sequence consistency with human Jun (also sharing a common ancestor with the Nach-CNC family) (Fig. S1B), whilst another original protein MEQ of BATF has emerged in *Gallid herpesvirus 2*, with a 60.82% BRLZ sequence identity with the BATF subfamily (Fig. S1A). Collectively, it is thus postulated that a common far-reaching origin of bZIP proteins will be identifiable from the most ancestral hosts throughout a faraway distance to *herpesvirus* resulting in the BATF-MEQ and Jun-CNC subfamilies. In this evolutionary process, the first round of expansion and diversification of bZIP proteins is inferable to have occurred in discrepant protozoan lineages insofar as to generate at least 8 subfamilies, and the ensuing second round of expansion and diversification of bZIP proteins is likely to have occurred in diverse metazoan lineages (except for vertebrates), before becoming gradual in distinct vertebrates, albeit the latter genes are allowed for the maximal expansion and diversification. In addition, a small bZIP factor encoded by human T-cell leukemia virus type1 (HTLV-1) (i.e. HBZ, acting as a transcription repressor of viral replication and proliferation) (45, 46) was herein redefined to contain double BR region and a unique LZ region, which comprises seven rounds of heptad repeats and shares a highly homology with MEQ, Nach1, Nach2, NF-E2 p45 and Nrf1γ (Fig. S2F).

Interestingly, a subgroup of bZIP transcription factors with a unique conserved CNC domain (Fig. 2) comprise NF-E2 p45 subunit, and related factors Nrf1 (also called NFE2L1, along with a short form LCR-F1 and a long form TCF11), Nrf2 and Nrf3, together with the repressors Bach1 and Bach2. Amongst vertebrates, these CNC-bZIP proteins are highly conserved with their founding member *Drosophila melanogaster* Cnc protein and *Caenorhabditis elegans* Skn-1, but none of their orthologues are identified in plants and fungi (47). These CNC proteins except Skn-1 act as obligate heterodimers with sMaf (48) or other bZIP proteins such as Jun (49), for DNA-binding to target genes involved in cytoprotection against oxidative and other stresses (50). However, it is regretful, in our present opinion, for a limitation to only tracing the origin of CNC-bZIP proteins back to arthropods, although the conserved CNC domain was first identified in the *Cnc* gene product from the *Drosophila melanogaster* (23). Fortunately, an ancestor subfamily of Nach1-8 with high homology with all CNC-bZIP factors are here identified to be present in Echinodermata, Mollusca, Actiniaria, Placozoa, Porifera and bacteria, respectively (Fig. 2). This discovery indicates that the CNC-bZIP proteins are originated from the marine bacteria to the primitive multicellular organism (e.g. *Amphimedon queeslandica*), the invertebrates and vertebrates, but seem to be also absent in the unicellular protozoans. It is important to note that only a single gene encoding the CNC-bZIP protein is found in each of those lower animals such as ascidians, sea urchin, octopus, fly and hydra, but *de facto* the expansion and diversification of distinct CNC-bZIP subfamily genes (each with different genomic loci in some species) are deduced to have occurred in the vertebrate evolutionary process.

There exists a specific subfamily of membrane-bound bZIP transcription factors that are folded into distinct topologies within and around the ER and then processed into a mature activator in order to be translocated into the nucleus before regulating cognate genes (Fig. 3). In the past two decades, the best understood membrane-bound transcription factors are a bZIP protein ATF6 and another bHLH-ZIP protein SREBP in mammals, which coordinately monitor expression of key genes responsible for biosynthesis of cholesterol and other lipids to meet cellular needs (51). Here, we have predicted by using two TM domains of SREBP as queries, and summarized all TM-containing bZIP proteins across 23 species, which were classified into 4 major subgroups of XBP1u (including the C-terminal TM region of Nrf1D), NHB1-CNC/Nach (i.e. NHB1-associated TM1 identified in Nrf1, Nrf3, CncC, Skn-1, as well as in newly identified Nach3, Nach5, Nach6 and Nach7), OASIS/ATF6 (with a single TM segment) and SREBPs. In response to ER stress, ATF6 and SREBP are allowed for a transfer from the ER to the Golgi apparatus, in which ATF6 and SREBP-TM1 are enabled for their successive proteolytic processing by SIP and S2P to yield their N-terminal releasable portions as activators (Fig. 3E), albeit a nuance in the processing of both proteins could determine their difference in the ER retention and release signals (52, 53). Amongst these bZIP proteins, the early presence of the TM domain is found in protozoans, but the TM-bound CNC-bZIP factors may be originated from the ancestral orthologue in Actiniaria rather than bacteria. However, none of the similar TM-containing bZIP proteins are searched in either unicellular organisms or prokaryotes (Fig. 3). It is hence inferable that such a TM-containing bZIP protein is likely encoded by a putative fusion gene, in which a small TM-encoding fragment was fused together with the original BRLZ-encoding gene locus during their biological evolutionary process. It is also plausible that the absence of TM-containing bZIP factors contributes to a simple biology process in unicellular organisms and prokaryotes. With the increasing complexity of biological behaviors along with distinct evolutionary morphologies, a naturally selected fusion of the TM region with the conserved BRLZ domain is presumed, in order to generate such a TM-fused bZIP transcription factor allowing it to be involved in signal transduction, ion transmission and other life processes. This is also further supported by the fact that the TM helices can be conferred on the fusion TM-bZIP proteins to play essential roles in distinct biological processes (54), as consistent with the notion that the TM-containing bZIP players regulate the ER functions in distinct UPR signaling to defense against the ER-derived stress (55).

Since the complex relationship between an organismic genotype and phenotype is clearly mediated by many of interrelated biochemical networks (56), Metazoans have evolutionarily developed a considerably higher proportion of heterodimeric bZIP interactions to homodimeric ones, along with more network complexity than those generated in the unicellular species (39). Herein, we have also demonstrated that the complex regulatory networks of human bZIP transcription factors, in which the CNC-bZIP factors are closely interactive with sMaf (i.e. MafG, MafF and MafK) (Fig. 4). Further examinations revealed that either knockout of Nrf1α or its constructive induction also enables it to trigger distinct and even opposing expression profiles of other bZIP genes. These certain genes only need to be active in a particular cell at any given time, but the transcriptional activity of such genes is finely monitored by upstream bZIP factors as a functional homo- or hetero-dimer for specifically binding to the genomic DNA motifs, such as AP1-like ARE sequences, in order to switch target genes on or off as required for stimulation or silence by distinctive biological cues. These bZIP transcription factors are also often working together to regulate basal and stimulated expression of some specific genes involved in the response to various intracellular and extracellular signals, and other stresses from the changing environments. Conversely, the failure of these bZIP factors to control the activity of given genes ultimately results in the pathogenesis of cancer, diabetes or a wide array of other diseases. Thus, the complex regulatory network of bZIP subfamily through interacting with their dimeric partners and/or other different members to control transcriptional expression of many genes is ensured to act as a key means for diverse cellular responses to complex and in constant environments, as described by authors referenced (57). Overall, diverse interactions of distinct bZIP factors with different partners elicit different regulatory effects on target genes. In turn, such these regulatory effects may also be monitored by their conserved functional motifs.

The present study demonstrates that these two closer Nach and CNC subfamilies of bZIP proteins with similar conserved but slightly different structural domains (e.g. NTD, Neh2L, Neh5L, NehL1 and Neh3L) and functional motifs (e.g. DLG, ETGE, DIDLID/DLG and DSGLSL) (Figs. 3 and 5, and also see Fig. S1) are rationally inferable to be originated from the marine bacterial orthologues (i.e. Nach1/2), which share a very early ancestral homologue with the Jun subfamily diversified from the original virus (at least herpesvirus, as we have identified herein). Further evidence has been also provided by us and other groups (58–60) has revealed that the canonical DSGxSL degron and another non-canonical DSGxxL motif within the Nach/CNC factors are involved in the regulation of their protein stability and transcriptional ability to mediate expression of AP1-like ARE-driven genes, responsible for antioxidant, detoxification and cytoprotection against cellular stress. Since NF-E2 p45 and Nrf3 are subject to their tissue-specific expression in haematopoietic and placental cell lineages respectively (61–63), transcriptional expression of such ARE-driven genes is therefore regulated primarily by two master players Nrf1 and Nrf2, essential for maintaining cellular (redox) homoeostasis and organ integrity in mammals. The activity of Nrf2 is negatively regulated by its DSGISL motif acting as a redox-insensitive β-TrCP (β-transducin repeat-containing protein)-binding degron (58, 59). Similar DSGLSL motif in Nrf1 was also identified as a GSK-3β-mediated phosphodegron targeting this CNC-bZIP to the β-TrCP^SCF^-dependent ubiquitin proteasomal degradation pathway (60). Excitingly, the activity of ARE-driven reporter gene regulated by Nrf1 and Nrf2 is significantly suppressed by their ancestral homologues Nach1 or Nach2 (both lack the Neh5L as an essential transactivation domain), and the suppression is also abolished by a mutant of Nach1 with a deletion of the canonical DSGxSL motif, but not by another point-mutant DAGxAL (Fig. 5). Further experiments determine that the DSGxSL motif also acts as a degron targeting Nach1 to the proteasomal degradation pathway, but the degradation is not completely prevented by a proteasomal inhibitor. Moreover, both the canonical DSGxSL and non-canonical DSGxxL degrons of related Nach/CNC-bZIP proteins appear to share a certain conservativity with the enzymatic active site (DSGxQx) of the DDI aspartic proteases (Fig. 5B). Collectively, it is thus reasonable that these Nach/CNC-bZIP proteins may also be auto-destructed by their DSGxSL or DSGxxL degrons *per se* (i.e. suicidon designated herein); this process enables target genes to be rapidly recovered after being switched off.

In conclusion, ever-accumulating databases of sequenced massive genomes from a large variety of species have led us to search for the primordial bZIP proteins before the ensuing reconstruction of their evolutionary trajectories within the neighbor-joining phylogenetic tree. Here, 495 of bZIP proteins were selected from 23 distinct genomes with representatives from metazoan, protozoan, bacteria to viruses, some of which are predicatively classified into four subgroups of the TM-bound bZIP subfamily (including NHB1-CNC/Nach, XBP1u, ATF6/OASIS, and SREBPs), in addition to an emphasis of Nrf1 involved in complex interaction networks of human bZIP proteins. Notably, two ancestral bZIP proteins of Jun and BATF subfamilies are originated within viruses, and the Jun subfamilies are evolutionarily conserved with another two closer Nach and CNC subfamilies that are diversified inferably from the marine bacteria orthologues. Thereafter, expansion and diversification of the bZIP superfamily are deduced to have occurred amongst disparate vertebrates from metazoan, whilst the TM-containing bZIP proteins should be originated from protozoans. More excitingly, we also discover an ancestral subfamily of Nach1-8 proteins, of which the marine bacteria Nach1/2 shares a graded homology with human NF-E2 p45, Nrf1 and Nrf2 of the CNC-bZIP subfamily, with distinct transcriptional abilities to mediate differential expression of ARE-battery genes.

## MATERIALS AND METHODS

### Identification of bZIP proteins

A BLAST programme was conducted to identify bZIP family members with the parameter (*E*-value = *e*^−5^). Meanwhile, another tool HMM (http://hmmer.org/) was also employed to identify bZIP proteins with the default parameter (*E*-value = 0.01). The resulting searched sequences were downloaded from the database of NCBI (National Center for Biotechnology), of which the repeated and incomplete sequences were manually removed. In addition, all the past not-yet-identified bZIP TFs were herein denoted by a nomenclature rule to give the acronym of species termed, i.e. xx_bZIP_TFx (Fig. S5). Notably, all the false-positive sequences were removed according to two selection criteria: Firstly, the clan CL0018 from the Pfam database contains three member families (PF00170, PF07716 and PF03131); secondly, all the BRLZ domains were identified by using the SMART sequence analysis.

### Phylogenic analysis of structural domains

For phylogenetic analysis of the BRLZ domains, they were extracted from all the selected bZIP proteins by using a local Perl script program, and then aligned using three tools DNAMAN 8.0, T-Coffee Server and ClustalX 2.0 with distinct default parameters. The multiple sequence alignments were manually refined and end-trimmed to eliminate the poor scored or divergent regions. Subsequently, the remaining unambiguously aligned sequences were subjected to construction of the neighbor-joining phylogenetic trees, by using the MEGA version 6.0 (with a gap treatment: partial deletion; model of evolution: the Poisson model; 1000 bootstrap replications), which are displayed by the iTOL programme (64). Moreover, the conservative motifs within CNC and BRLZ domains were also analyzed by the MEME and Web-logo tools with different parameters. In addition, the secondary structures of CNC and adjacent BRLZ domains were predicted by using the PSIPRED tool and the 3D structures of Nrf1 and Nach1 with the homology of Skn-1 were further modeled by the SWISS-MODEL software.

### Bioinformatic analysis of TM-containing transcription factors

All of the above-selected bZIP proteins were also redefined so as to discover putative TM-containing transcription factors by using two different programs called TMpred and TMHMM. These TM domains were then subjected to the phylogenetic analysis, with multiple sequence alignment and conservative analysis. In addition, six TM-folded α-helix properties of Nrf1, XBP1u, ATF6, OASIS, SREBP-TM1 and -TM2 were calculated with the HeliQuest tool.

### Interaction network and transcriptomic analysis

The interaction networks of between bZIP proteins in humans were constructed with the STRING software (a <u>*S*</u>earch <u>*T*</u>ool for the <u>*R*</u>etrieval of *I*nteracting <u>*G*</u>enes and/or proteins, http://string-db.org/) (65). Relative levels of gene expression were calculated as RPKM (<u>*R*</u>eads <u>*P*</u>er <u>*K*</u>ilobase per <u>*M*</u>illion mapped reads). According to the Log2-based RPKM value, the heat map was also generated with the MEV4.9 program.

### Experimental cell lines

Experimental cell lines, including human hepatocellular carcinoma HepG2 (i.e. *Nrf1^+/+^*), *Nrf1α^−/−^* (established by Talens-mediated *Nrf1α*-specific knockout in HepG2), human embryonic kidney (HEK293) and HEK293C^Nrf1α^ (with stable tetracycline-inducible expression of Nrf1α established in human embryonic kidney) were cultured in a 37℃ incubator with 5% carbon dioxide, and allowed for growth in Dulbecco’s modified Eagle’s medium (DMEM) with 25 mmol/L high glucose, 10% (v/v) fetal bovine serum (FBS), 100 units/ml penicillin-streptomycin.

### Validation of gene expression by qRT-PCR

Total RNAs was extracted from cell samples, by using an RNA extraction kit (TIANGEN, China) and then subjected to reactions with reverse transcriptase (Promega, USA) to synthesize the single-strand cDNAs. Subsequently, expression of the indicated bZIP genes at mRNA levels in different cell lines were measured by qRT-PCR, which was carried out in the GoTaq® real-time PCR detection systems, loaded on a CFX96 instrument (Bio-rad, USA). The results were analyzed by the Bio-Rad CFX Manager 3.0 software.

### The pulse-chase experiments followed by Western blotting

After reaching 70% confluence of HepG2 cells that had been allowed for growth in 6-well plates for 24 h in DMEM contain 25 mmol/L glucose and 10% FBS, they were transfected with an expression construct for Nrf1, Nrf2, Nach2, Nach1 or its mutants (all were tagged C-terminally by the V5 epitope) in a mixture of Lipofectamine 3000 (Invitrogen). After transfection for 24 hours, the cells were or were not treated with CHX (at 50 µg/ml) alone or plus MG132 (at 10 µmol/L) for additional 30 min to 4 h before being harvested in a lysis buffer (36). Total cell lysates were subjected to protein separation by SDS-PAGE gels containing 10% polyacrylamide, followed by Western blotting with antibodies against the V5 epitope (Invitrogen Ltd) or β-Actin (from ZhongshanJinqiao Co, Beijing, China). β-Actin served as an internal control to verify amounts of proteins that were loaded in each well occasion. The intensity of Nach1 and its mutant protein bands was quantified and shown graphically.

### ARE-driven reporter gene assays

Equal numbers (1.5×10^5^) of HepG2 cells were allowed for 24-h growth in each well of 12-well plates containing DMEM supplemented with 25 mmol/L glucose and 10% FBS. After reaching 70% confluence, the cells were transfected with expression constructs for Nrf1, Nrf2, Nach2, Nach1 or its mutants alone or in different combinations with one another, together with both plasmids of *GSTA2-6×ARE-Luc* reporter and pRL-TK (as an internal control), in a mixture of Lipofectamine 3000 (Invitrogen). Approximately 24 h after transfection, ARE-driven luciferase reporter activity was measured by Magellan7.1 SP1 systems and then calculated as fold changes (mean ± S.D), as described previously (66). The data presented each represent at least three independent experiments.

### Statistical analysis

Statistical significances of fold changes in the *GSTA2-6×ARE-Luc* reporter activity and also in the gene expression were determined using the *Student’s* t-test or Multiple Analysis of Variations (MANOVA). The data are shown as a fold change (mean ± S.D), each of which represents at least 3 independent experiments that were each performed in triplicates.

## Acknowledgments

We are greatly thankful to Prof. Ze Zhang for his expertise in molecular evolution. The study was supported by the National Natural Science Foundation of China (key programs 91129703, 91429305 and project 31270879) awarded to Prof. Yiguo Zhang (University of Chongqing, China), and in part funded by the Chongqing University postgraduates innovation project (No. CYB15024) awarded to Mr. Lu Qiu.

## Author contributions

Y.Z. performed all bioinformatics analyses and most experiments, collected the resulting data and prepared drafts of this manuscript with most figures. M.W. helped Y.Z. with the interactive networks of human bZIP proteins and validation of gene expression by RT-qPCR. Y.X. performed Western blotting of Nach1 and Mut1 with distinct half-lives estimated. L.Q. helped Y.Z. with luciferase reporter assays. S.H. helped Y.Z. together with molecular cloning to create expression constructs. P.M. and Z.Z. contributed to critical scientific discussions and editorial skills to revise the manuscript. Lastly, YG.Z. designed this study, analyzed all the data, helped to prepare all figures, wrote and revised the paper.

## Data and materials availability

All data needed to evaluate the conclusions in the paper are present in the paper and/or the Supplementary Materials. Additional data related to this paper may be requested from the authors.

## SUPPLEMENTARY MATERIALS

**Fig. S1. Identity of original homologues of BATF, Jun and C/EBP subfamilies.**

Multiple sequence alignments of BRLZ domains were analyzed by the DNAMAN8.0 software: (**A**) MEQ from *Gallid herpesvirus 2* is evolutionarily closer to human BATF family; (**B**) Another viral original protein (with a GenBank No. YP_007003813) from *Cyprinid herpesvirus 1* is conserved with human Jun family; (**C**) An additional bacterial homologous protein (with a GenBank accession No. WP_062270874) from *Endozoicomonas arenosclerae* is classified into human C/EBP family.

**Fig. S2. Detailed schematic representation of structural domains of Nach/CNC-bZIP proteins**.

Bioinformatic analysis by the DNAMAN8.0 software was subject to multiple sequence alignments of different structural domains of: (**A**) the Neh2L domain, (**B**) the Neh5L domain, (**C**) Neh3L domain. (**D, E**) Nach1 and Nach2 have no effects on basal expression of AP-1-driven reporter gene and its regulation by Fos and Jun. Related methods and data calculations were referenced to determination of ARE-driven luciferase reporter activity as described in the legend of main Fig. 5. (**F**) Shows specific sequence alignments of the BRLZ domains of HBZ from *Human T-Cell Leukemia Virus Type 1*, MEQ from *Gallid herpesvirus 2*, bacterial Nach1/2 with human Nrf1γ. (**G**) Shows an additional alignment of the full length Nach1/2 proteins with human NF-E2 P45.

**Fig. S3~S13. Distinct characteristics of BRLZ domains within different bZIP subfamilies**.

Distinct characteristics of BRLZ domain were analyzed by using three different softwares DNAMAN8.0, MEME and Web-logo with default parameters.

**Fig. S3**. Alignment of the CNC domains from those identified Nach/CNC-bZIP subfamily proteins.

**Fig. S4**. Alignment of the BRLZ domains from those identified Nach/CNC-bZIP subfamily proteins.

**Fig. S5**. Alignment of the BRLZ domains of unclassified bZIP proteins with human bZIP representatives.

**Fig. S6**. Alignment of the BRLZ domains from within both Maf and sMaf subfamilies.

**Fig. S7.** Alignment of the BRLZ domains from within both Fos (**A**) and Jun (**B**) subfamilies.

**Fig. S8.** Alignment of the BRLZ domains from within both ATF6 (**A**) and OASIS (**B**) subfamilies.

**Fig. S9.** Alignment of the BRLZ domains from within both ATF2 (**A**) and ATF4 (**B**) subfamilies.

**Fig. S10.** Alignment of the BRLZ domains from within both ATF3 (**A**) and BATF (**B**) subfamilies.

**Fig. S11.** Alignment of the BRLZ domains from within both PAR (**A**) and E4BP4 (**B**) subfamilies.

**Fig. S12.** Alignment of the BRLZ domains from within the C/EBP subfamilies.

**Fig. S13.** Alignment of the BRLZ domains from within both CREB (**A**) and XBP1 (**B**) subfamilies.

**Fig. S14. The whole images of figure 5I**.

(**A** to **C**) Western blotting of Nach1 and its Mut1 that had been resolved by the whole PAGE gels containing 10% polyacrylamide, of which the cropped images were also shown in the main I1, I2 and I3 in Fig. 5I, respectively.

