## Supplementary Materials for "Nach is a novel ancestral subfamily ofthe CNC-bZIP transcription factors selected during evolution from the marine bacteria to human"

**fig. S1. Identity of original homologues of BATF, Jun and C/EBP subfamilies.**

Multiple sequence alignments of BRLZ domains were analyzed by the DNAMAN8.0 software: (**A**) MEQ from *Gallid herpesvirus 2* is evolutionarily closer to human BATF family; (**B**) Another viral original protein (with a GenBank No. YP_007003813) from *Cyprinid herpesvirus 1* is conserved with human Jun family; (**C**) An additional bacterial homologous protein (with a GenBank accession No. WP_062270874) from *Endozoicomonas arenosclerae* is classified into human C/EBP family.

**fig. S2. Detailed schematic representation of structural domains of Nach/CNC-bZIP proteins.**

Bioinformatic analysis by the DNAMAN8.0 software was subject to multiple sequence alignments of different structural domains of: (**A**) the Neh2L domain, (**B**) the Neh5L domain, (**C**) Neh3L domain. (**D, E**) Nach1 and Nach2 have no effects on basal expression of AP-1-driven reporter gene and its regulation by Fos and Jun. Related methods and data calculations were referenced to determination of ARE-driven luciferase reporter activity as described in the legend of main Fig. 5. (**F**) Shows specific sequence alignments of the BRLZ domains of HBZ from *Human T-Cell Leukemia Virus Type 1*, MEQ from *Gallid herpesvirus 2,* bacterial Nach1/2 with human Nrf1γ. (**G**) Shows an additional alignment of the full length Nach1/2 proteins with human NF-E2 P45.

**fig. S3~S13. Distinct characteristics of BRLZ domains within different bZIP subfamilies.**

Distinct characteristics of BRLZ domain were analyzed by using three different softwares DNAMAN8.0, MEME and Web-logo with default parameters.

**fig. S3**. Alignment of the CNC domains from those identified Nach/CNC-bZIP subfamily proteins.

**fig. S4**. Alignment of the BRLZ domains from those identified Nach/CNC-bZIP subfamily proteins.

**fig. S5**. Alignment of the BRLZ domains of unclassified bZIP proteins with human bZIP representatives.

**fig. S6**. Alignment of the BRLZ domains from within both Maf and sMaf subfamilies.

**fig. S7.** Alignment of the BRLZ domains from within both Fos (**A**) and Jun (**B**) subfamilies.

**fig. S8.** Alignment of the BRLZ domains from within both ATF6 (**A**) and OASIS (**B**) subfamilies.

**fig. S9.** Alignment of the BRLZ domains from within both ATF2 (**A**) and ATF4 (**B**) subfamilies.

**fig. S10.** Alignment of the BRLZ domains from within both ATF3 (**A**) and BATF (**B**) subfamilies.

**fig. S11.** Alignment of the BRLZ domains from within both PAR (**A**) and E4BP4 (**B**) subfamilies.

**fig. S12.** Alignment of the BRLZ domains from within the C/EBP subfamilies.

**fig. S13.** Alignment of the BRLZ domains from within both CREB (**A**) and XBP1 (**B**) subfamilies.

**fig. S14. The whole images of figure 5I.**

(**A** to **C**) Western blotting of Nach1 and its Mut1 that had been resolved by the whole PAGE gels containing 10% polyacrylamide, of which the cropped images were also shown in the main I1, I2 and I3 in Fig. 5I, respectively.

fig. S1

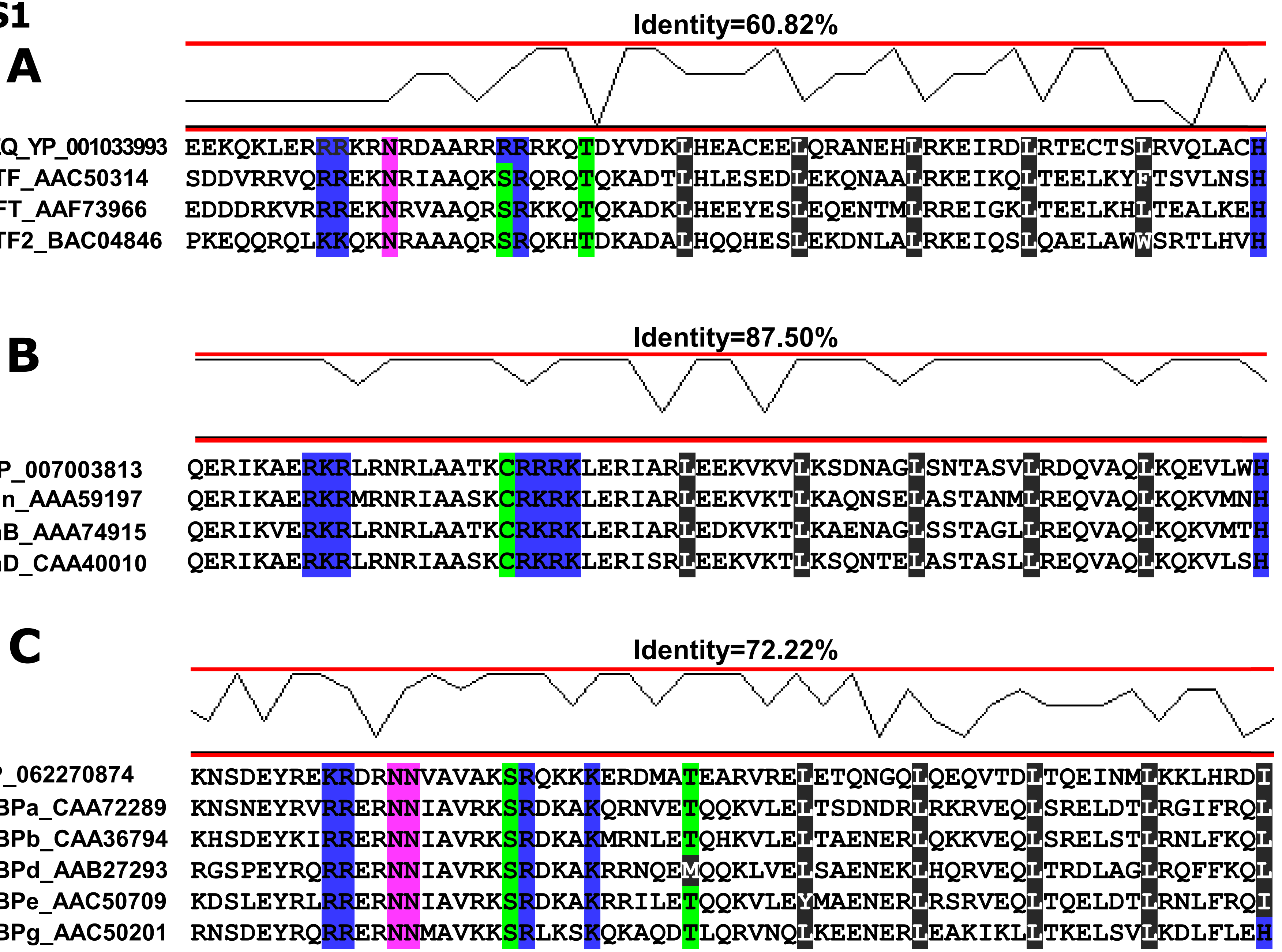
A
Identity=60.82%
Gh2-MEQ_YP_001033993 EEKQKLERRRKRNRDAARRRRRKQTDYVDKLHEACEELQRANEHLRKEIRDLRTECTSLRVQLACH
Hs-BATF_AAC50314 SDDVRRVQRREKNRIAAQKSRQRQTQKADTLHLESEDLEKQNAALRKEIKQLTEELKYFTSVLNSH
Hs-SNFT_AAF73966 EDDDRKVRRREKNRVAAQRSRKKQTQKADKLHEEYESLEQENTMLRREIGKLTEELKHLTEALKEH
Hs-BATF2_BAC04846 PKEQQRQLKKQKNRAAAQRSRQKHTDKADALHQQHESLEKDNLALRKEIQSLQAELAWWSRTLHVH
B
Identity=87.50%
Ch1_YP_007003813 QERIKAERKRLRNRLAATKCRRRKLERIARLEEKVKVLKSDNAGLSNTASVLRDQVAQLKQEVLWH
Hs-cJun_AAA59197 QERIKAERKRMRNRIAASKCRKRKLERIARLEEKVKTLKAQNSELASTANMLREQVAQLKQKVMNH
Hs-JunB_AAA74915 QERIKVERKRLRNRLAATKCRKRKLERIARLEDKVKTLKAENAGLSSTAGLLREQVAQLKQKVMTH
Hs-JunD_CAA40010 QERIKAERKRLRNRIAASKCRKRKLERISRLEEKVKTLKSQNTELASTASLLREQVAQLKQKVLSH
C
Identity=72.22%
Ea_WP_062270874 KNSDEYREKRDRNNVAVAKSRQKKKERDMATEARVRELETQNGQLQEQVTDLTQEINMLKKLHRDI
Hs-CEBPa_CAA72289 KNSNEYRVRRERNNIAVRKSRDKAKQRNVETQQKVLELTSDNDRLRKRVEQLSRELDTLRGIFRQL
Hs-CEBPb_CAA36794 KHSDEYKIRRERNNIAVRKSRDKAKMRNLETQHKVLELTAENERLQKKVEQLSRELSTLRNLFKQL
Hs-CEBPd_AAB27293 RGSPEYRQRRERNNIAVRKSRDKAKRRNQEMQQKLVELSAENEKLHQRVEQLTRDLAGLRQFFKQL
Hs-CEBPe_AAC50709 KDSLEYRLRRERNNIAVRKSRDKAKRRILETQQKVLEYMAENERLRSRVEQLTQELDTLRNLFRQI
Hs-CEBPg_AAC50201 RNSDEYRQRRERNNMAVKKSRLKSKQKAQDTLQRVNQLKEENERLEAKIKLLTKELSVLKDLFLEH

fig. S2

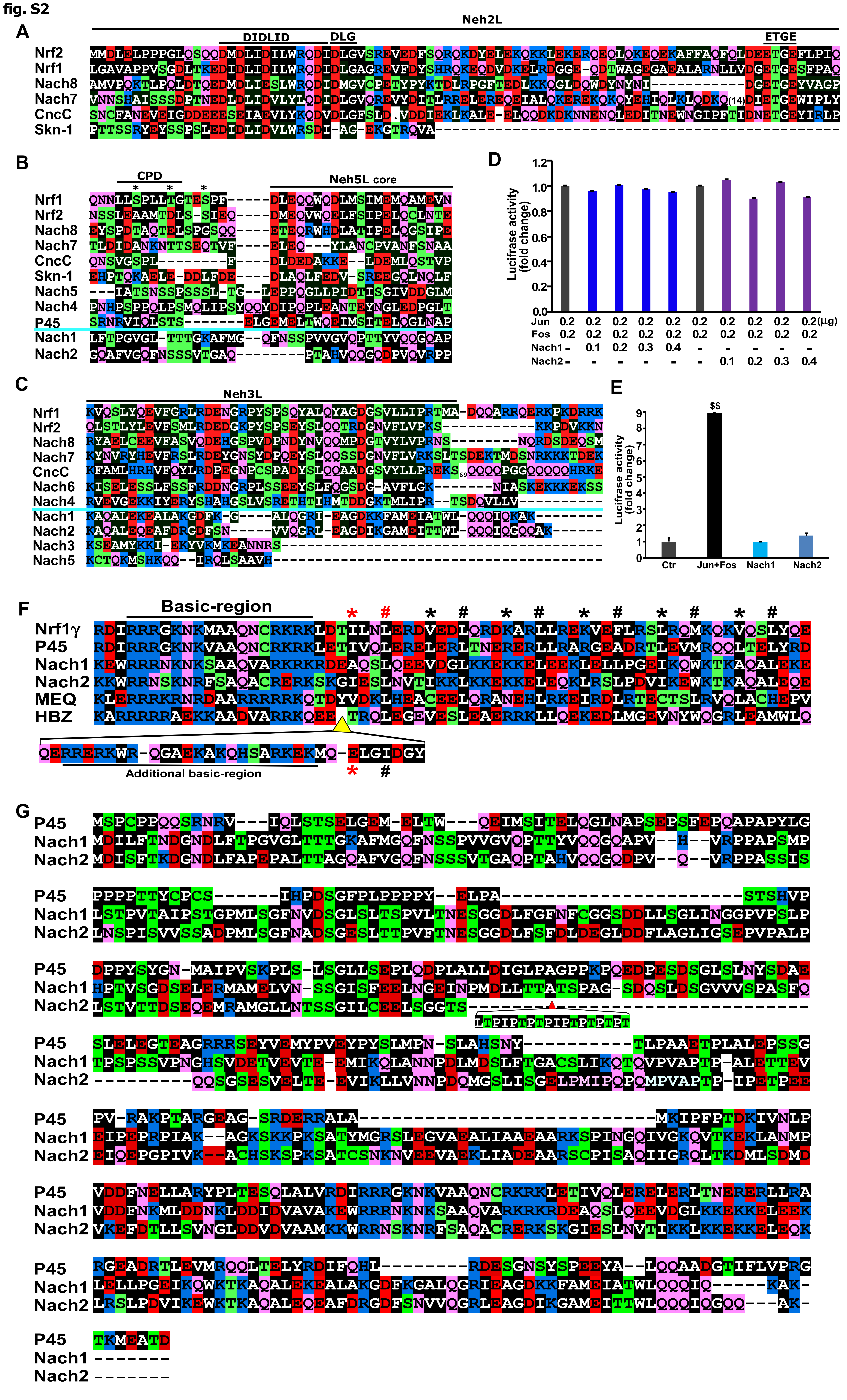
A
Neh2L
DIDLID
DLG
ETGE
Nrf2
Nrf1
Nach8
Nach7
CncC
Skn-1
B
CPD
Neh5L core
Nrf1
Nrf2
Nach8
Nach7
CncC
Skn-1
Nach5
Nach4
P45
Nach1
Nach2
C
Neh3L
Nrf1
Nrf2
Nach8
Nach7
CncC
Nach6
Nach4
Nach1
Nach2
Nach3
Nach5
D
Luciferase activity (fold change)
Jun 0.2 0.2 0.2 0.2 0.2 0.2 0.2 0.2 0.2 0.2(μg)
Fos 0.2 0.2 0.2 0.2 0.2 0.2 0.2 0.2 0.2 0.2
Nach1 - 0.1 0.2 0.3 0.4 - - - - -
Nach2 - - - - - - 0.1 0.2 0.3 0.4
E
Luciferase activity (fold change)
$$
Ctr
Jun+Fos
Nach1
Nach2
F
Basic-region
Nrf1γ
P45
Nach1
Nach2
MEQ
HBZ
Additional basic-region
G
P45
Nach1
Nach2

fig. S3

### CNC-bZIP

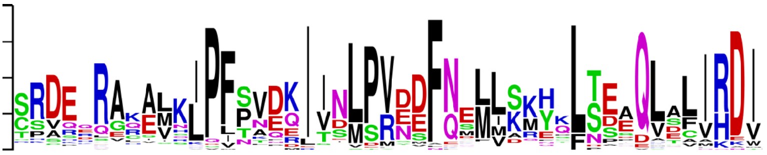

| Group | Sequence | Alignment |
|---|---|---|
| Bach1 | Hs-Bach1_NP_001177 | CSAREQECEVKLPFNAQRIISLSRNDFQSLLKMHKLTPEQLDCIHDI |
| | Gg-Bach1_XP_416696 | CSAREQECEVKLPFNAQRIISLSRNDFQSFLKMHKLTPEQLDCIHDI |
| | Mm-Bach1_NP_031546 | CSAREQDCEVKLPFNAQRIISLSRNDFQSLLKMHKLTPEQLDCIHDI |
| | Ac-Bach1_XP_003219110 | CSAREQECEIKLPMNPQKIISLSRNDFQSFLKMHKLTPEELDCIHDI |
| | Xl-Bach1a_XP_018101443 | ESCSVQECETKLPFNAQKIISLSRYDFQALVKKHSLSAEQLDCIHDI |
| | Xl-Bach1b_XP_018104014 | CSAKELECEEKLPFKAQKIISLSRYDFQALIKRHSLTAEQLDCIHDI |
| Bach2 | Hs-Bach2_NP_068585 | CPVQDRGQEVKLPFPVDQITDLPRNDFQMMIKMHKLTSEQLEFIHDV |
| | Mm-Bach2_NP_001103131 | CPVQDRGQEVKLPFPVDQITDLPRNDFQMMIKMHKLTSEQLEFIHDI |
| | Ac-Bach2_XP_016852773 | CPVQDRGHEVKLPFPVDQITDLPRNDFQMMVKMHKLTSEQLEFIHDV |
| | Gg-Bach2_XP_015140173 | CPVQDRGHEVKLPFPVDQITDLPRNDFQMMIKMHKLTSEQLEFIHDV |
| | Xl-Bach2a_XP_018118876 | CPVQDRGHEVKLPFPVDQITDLPRNDFQMMIKMHKLSSEQLEFIHDV |
| | Xl-Bach2b_XP_018120920 | CPVQDSGQEVKLPFPVDQITDLPRNDFQMMVKMHKLSSEQLEFIHDV |
| | Dr-Bach2b_XP_005160605 | CHVREQGPEVKLPFPVDQITNLPRNDFQMMVKMHKLTSEQLEFIHDV |
| | Dr-Bach2a_XP_685315 | FATKERTQEVKLPFSVDQITELPRNDFQLMIKMHKLSSDQLDFIHDM |
| | *Dr-Bach1b_AHX22538 | SYSSERVREMALPLSVDQIVCLSRNDFQQMLKQQSLSREQLDAVHDI |
| Nrf3 | Hs-Nrf3_NP_004280 | SRDEQRAKALHIPFSVDEIVGMPVDSFNSMLSRYYLTDLQVSLIRDI |
| | Mm-Nrf3_NP_035033 | SRDEQRAKALHIPFSVDEIVRMPVDSFNSMLSRYYLTDLQVSLIRDI |
| | Xl-Nrf3_XP_018122887 | SRDERRAKSLRIPFSVDEIVSMPVEMFNNMLSKHCLTESQVTVIRDI |
| | Xl-Nrf3L_XP_018124724 | SRDERRAKSLRIPFSVDEIVSMPVEMFNNMLSKHCLTESQVTVIRDI |
| | Ac-Nrf3_XP_003222201 | SHDEYCVKALRIPFAINDIVTLPVDSFNSMLSKYCLTDNQLSLIRDI |
| Nrf1 | Hs-Nrf1_NM_003204.2 | SRDEHRARAMKIPFTNDKIINLPVEEFNELLSKYQLSEAQLSLIRDI |
| | Mm-Nrf1_NM_008686.3 | SRDEHRARAMKIPFTNDKIINLPVEEFNELLSKYQLSEAQLSLIRDI |
| | Gg-Nrf1_NM_001030756 | SRDEHRARAMKIPFTNDKIINLPVEEFNELLSKYQLSEAQLSLIRDI |
| | Ac-Nrf1_XM_016994618 | SRDEHRARAMKIPFSNEKIINLPVEEFNELLSKYQLSEAQLSLIRDI |
| | Xl-Nrf1L2_NP_001080013 | GRDEQRAKAMKIPFTNDKIINLPVEEFNDLLAKYQLSEAQLCLIRDI |
| | Xl-Nrf1L1_XP_018090244 | GRDEQRARAMKIPFSNDKIINLPVEEFNDLLAKYQLSEAQLCLIRDI |
| | Dr-Nrf1b_NP_001265771 | SRDERRARAMDIPFSNEKIINLPVEEFNELLAKHHLNEDQLSLIRDI |
| | Dr-Nrf1a_JX867114 | SRDEKRARLMNIPFSNERIINLTVDEFNRLLEKYHLSEAQSHANQNT |
| Nrf2 | Hs-Nrf2_NP_006155 | TRDELRAKALHIPFPVEKIINLPVVDFNEMMSKEQFNEAQLALIRDI |
| | Mm-Nrf2_NP_035032 | TRDELRAKALHIPFPVEKIINLPVDDFNEMMSKEQFNEAQLALIRDI |
| | Ac-Nrf2_XP_003226813 | TRDELRAKALQIPFPVEKIINLPVDDFNEMMSKEQFTEAQVTLIRDI |
| | Gg-Nrf2_NP_990448 | TRDEQRAKALQIPFPVEKIINLPVDDFNEMMSKEQFSEAQLALIRDI |
| | Dr-Nrf2b_NP_001244112 | SRDEQRAKALQIPFTVDMIINLPVDDFNEMMSKHQLNEAQLALVRDI |
| | Xl-Nrf2L_NP_001079522 | TRDEQRAKVLNLPFTVEKIVNLPVDSFNEIMSKYQFNEAQLALIRDI |
| | *Dr-Nrf3_NP_998396 | SRDEHRVQALGLPFSAFQIVNMPVENFLELLDRQNLSGSDVTLLRDV |
| | Dr-Nrf2a_NP_878309 | CRDEQRAQALSLPLSVHDIIHLPVEAFNEAISTCKLNHAQHTLIRDI |
| | Xl-Nrf2S_NP_001086307 | TRDTQRAKVLNIPFSVDKIVNLPVDSFNEMMSKYQFNEAQLALIRDI |
| p45 | Hs-P45_NP_006154 | SRDERRALAMKIPFPTDKIVNLPVDDFNELLARYPLTESQLALVRDI |
| | Mm-p45_NP_032711 | SRDERRALAMKIPFPTDKIVNLPVDDFNELLAQYPLTESQLALVRDI |
| | Xl-p45_XP_018103033 | SRDERRAAAMNIPFPTERIVNLPVEDFNELLSRYTLTDTQLALVRDI |
| | Dr-P45_NP_778208 | SRDERRALALKIPFSLEKIVNLPVDDFNELLTQFTLTDAQLALVRDI |
| | Ac-P45_XP_003216776 | SRDERRALAMKIPFPVEKIINLPVDDFNELVSRFPLSEPQLALIRDI |
| Nach | *Dm-CncC_NP_732833 | TRDEKRARSLNIPISVPDIINLPMDEFNERLSKYDLSENQLSLIRDI |
| | Nv-Nach6_AOP32002 | LASKADADSLDIGVSEEKIVEMSVAEFTTFLEK..LSDAQAKYVRDV |
| | *Ta-Nach3_XP_002116568 | IFGVPTNSMLVLPVPERELVDMPVNEFLAMIER..LPSDVAALARDV |
| | *Sp-Nach8_NP_001123278 | NRDEKRAKALKLPVCLEKIINLPVDSFNDLVKKYELTDPQMQLVRDI |
| | *Ob-Nach7_XP_014784776 | SRDHKRLQELKVPLSMNQIINSSVEEFNDLVRKYPLPEPQLRLIRDI |
| | Aq-Nach5_XP_003385261 | SLASPDIPQSFNGISKLEIIDMPYFEFKKLLDGGTFSERDKEEVKAI |
| | *Aq-Nach4_XP_011402550 | MSSHNRRGGASKPITRDKLVIMPVEDFNSLLDEALLSEIEVAFMKEW |
| | Es-Nach2_WP_062268396 | RKSPINGQIVGKQVTKEKLANMPVDDFNKMLDDNKLDDIDVAVAKEW |
| | En-Nach1_WP_034835593 | RSCPISAQIIGRQLTKDMLSDMDVKEFDTLLSVNGLDDVDVAAMKKW |
| | *Ce-Skn-1_NP_741404 | SKDEQLASDNELPVSAFQISEMSLSELQQVLKNESLSEYQRQLIRKI |
| | Dr-Bach1a_NP_001035403 | .......................MTRRAFLQMLKREQLTPEQLEYVQDV |

**fig. S4**

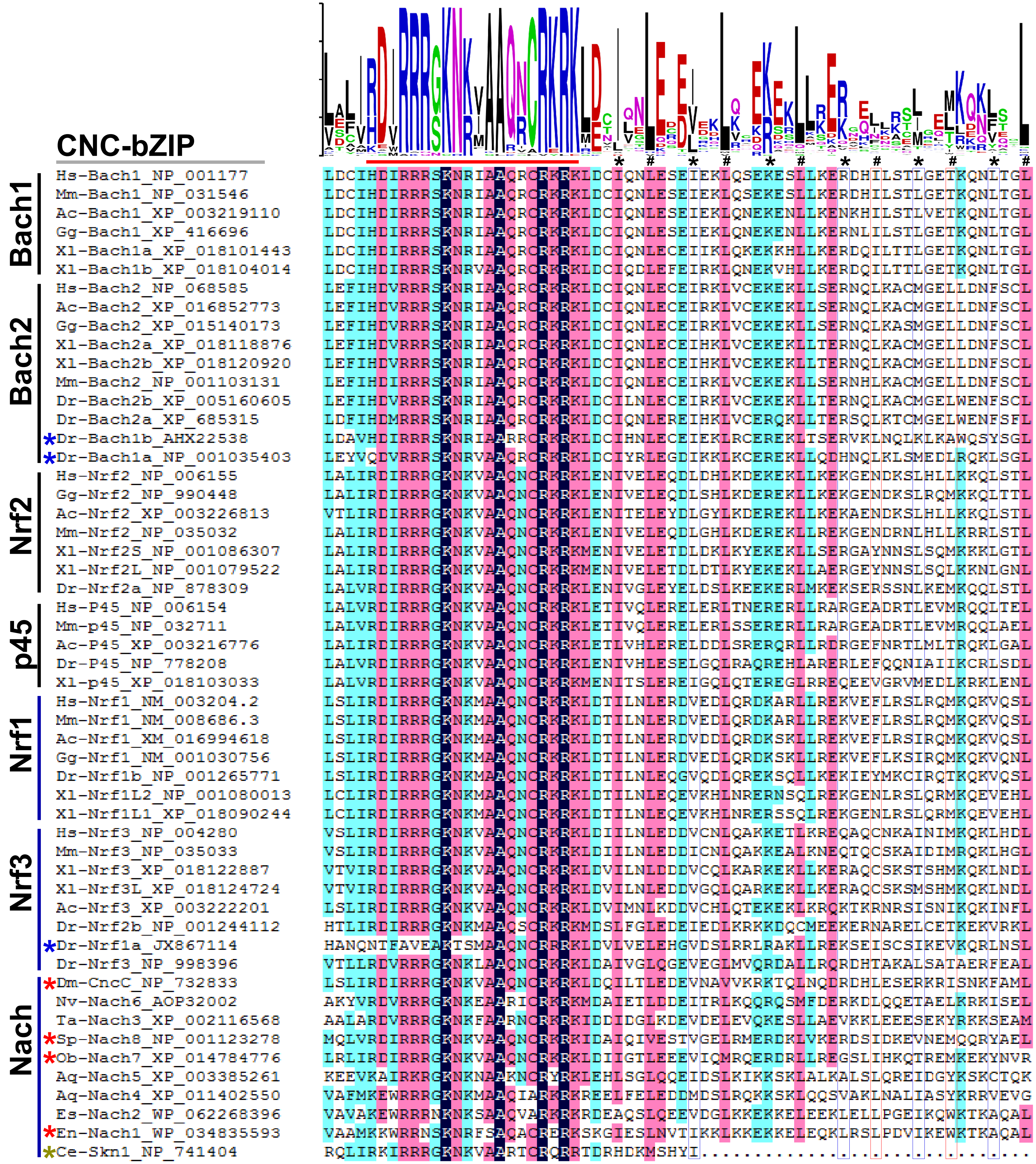
CNC-bZIP
Bach1
Bach2
Nrf2
p45
Nrf1
Nrf3
Nach
Hs-Bach1_NP_001177 LDCIHDIRRRSKNRIAAQRCRKRKLDCIQNLESEIEKLQSEKESLLKERDHILSTLGETKQNLTGL
Mm-Bach1_NP_031546 LDCIHDIRRRSKNRIAAQRCRKRKLDCIQNLESEIEKLQSEKESLLKERDHILSTLGETKQNLTGL
Ac-Bach1_XP_003219110 LDCIHDIRRRSKNRIAAQRCRKRKLDCIQNLESEIEKLQNEKENLLKENKHILSTLVETKQNLTGL
Gg-Bach1_XP_416696 LDCIHDIRRRSKNRIAAQRCRKRKLDCIQNLESEIEKLQNEKENLLKERNLILSTLGETKQNLTGL
Xl-Bach1a_XP_018101443 LDCIHDIRRRSKNRIAAQRCRKRKLDCIQNLECEIIKLQKEKKHLLKERDQILTTLGETKQNLTGL
Xl-Bach1b_XP_018104014 LDCIHDIRRRSKNRVAAQRCRKRKLDCIQDLEFEIRKLQNEKVHLLKERDQILTTLGETKQNLTGL
Hs-Bach2_NP_068585 LEFIHDVRRRSKNRIAAQRCRKRKLDCIQNLECEIRKLVCEKEKLLSERNQLKACMGELLDNFSCL
Ac-Bach2_XP_016852773 LEFIHDVRRRSKNRIAAQRCRKRKLDCIQNLECEIRKLVCEKEKLLSERNQLKACMGELLDNFSCL
Gg-Bach2_XP_015140173 LEFIHDVRRRSKNRIAAQRCRKRKLDCIQNLECEIRKLVCEKEKLLSERNQLKASMGELLDNFSCL
Xl-Bach2a_XP_018118876 LEFIHDVRRRSKNRIAAQRCRKRKLDCIQNLECEIHKLVCEKEKLLTERNQLKACMGELLDNFSCL
Xl-Bach2b_XP_018120920 LEFIHDVRRRSKNRIAAQRCRKRKLDCIQNLECEIHKLVCEKEKLLTERNQLKACMGELLDNFSCL
Mm-Bach2_NP_001103131 LEFIHDIRRRSKNRIAAQRCRKRKLDCIQNLECEIRKLVCEKEKLLSERNHLKACMGELLDNFSCL
Dr-Bach2b_XP_005160605 LEFIHDVRRRSKNRIAAQRCRKRKLDCILNLECEIRKLVCEKEKLLTERNQLKACMGELWENFSCL
Dr-Bach2a_XP_685315 LDFIHDMRRRSKNRIAAQRCRKRKLDCIQNLEREIHKLVCERQKLLTERSQLKTCMGELWENFSFL
*Dr-Bach1b_AHX22538 LDAVHDIRRRSKNRIAARRCRKRKLDCIHNLECEIEKLRCEREKLTSERVKLNQLKLKAWQSYSGL
*Dr-Bach1a_NP_001035403 LEYVQDVRRRSKNRVAAQRCRKRKLDCIYRLEGDIKKLKCEREKLLQDHNQLKLSMEDLRQKLSGL
Hs-Nrf2_NP_006155 LALIRDIRRRGKNKVAAQNCRKRKLENIVELEQDLDHLKDEKEKLLKEKGENDKSLHLLKKQLSTL
Gg-Nrf2_NP_990448 LALIRDIRRRGKNKVAAQNCRKRKLENIVELEQDLSHLKDEREKLLKEKGENDKSLRQMKKQLTTL
Ac-Nrf2_XP_003226813 VTLIRDIRRRGKNKVAAQNCRKRKLENITELEYDLGYLKDEREKLLKEKAENDKSLHLLKKQLSTL
Mm-Nrf2_NP_035032 LALIRDIRRRGKNKVAAQNCRKRKLENIVELEQDLGHLKDEREKLLREKGENDRNLHLLKRRLSTL
Xl-Nrf2S_NP_001086307 LALIRDIRRRGKNKVAAQNCRKRKMENIVELETDLDKLKYEKEKLLSERGAYNNSLSQMKKKLGTL
Xl-Nrf2L_NP_001079522 LALIRDIRRRGKNKVAAQNCRKRKMENIVELETDLDTLKYEKEKLLAERGEYNNSLSQLKKNLGNL
Dr-Nrf2a_NP_878309 LALVRDIRRRGKNKVAAQNCRKRKLENIVGLEYELDSLKEEKERLMKEKSERSSNLKEMKQQLSTL
Hs-P45_NP_006154 LALVRDIRRRGKNKVAAQNCRKRKLETIVQLERELERLTNERERLLRARGEADRTLEVMRQQLTEL
Mm-p45_NP_032711 LALVRDIRRRGKNKVAAQNCRKRKLETIVQLERELERLSSERERLLRARGEADRTLEVMRQQLAEL
Ac-P45_XP_003216776 LALIRDIRRRGKNKVAAQNCRKRKLETLVHLERELDDLSRERQRLLRDRGEFNRTLMLTRQKLGAL
Dr-P45_NP_778208 LALVRDIRRRGKNKVAAQNCRKRKLENIVHLESELGQLRAQREHLARERLEFQQNIAIIKCRLSDL
Xl-p45_XP_018103033 LALVRDIRRRGKNKVAAQNCRKRKMENITSLEREIGQLCTEREGLRREQEEVGRVMEDLKRKLENL
Hs-Nrf1_NM_003204.2 LSLIRDIRRRGKNKMAAQNCRKRKLDTILNLERDVEDLQRDKARLLREKVEFLRSLRQMKQKVQSL
Mm-Nrf1_NM_008686.3 LSLIRDIRRRGKNKMAAQNCRKRKLDTILNLERDVEDLQRDKARLLREKVEFLRSLRQMKQKVQSL
Ac-Nrf1_XM_016994618 LSLIRDIRRRGKNKMAAQNCRKRKLDTILNLERDVDDLQRDKSKLLREKVEFLRSIRQMKQKVQSL
Gg-Nrf1_NM_001030756 LSLIRDIRRRGKNKMAAQNCRKRKLDTILNLERDVEDLQRDKSKLLREKVEFLKSIRQMKQKVQNL
Dr-Nrf1b_NP_001265771 LSLIRDIRRRGKNKMAAQNCRKRKLDTILNLEQGVQDLQREKSQLLKEKIEYMKCIRQTKQKVQSL
Xl-Nrf1L2_NP_001080013 LCLIRDIRRRGKNKMAAQNCRKRKLDTILNLEQEVKHLNRERNSQLREKGENLRSLQRMKQEVEHL
Xl-Nrf1L1_XP_018090244 LCLIRDIRRRGKNKMAAQNCRKRKLDTILNLEQEVKHLNRERSSQLREKGENLRSLQRMKQEVEHL
Hs-Nrf3_NP_004280 VSLIRDIRRRGKNKVAAQNCRKRKLDIILNLEDDVCNLQAKKETLKREQAQCNKAINIMKQKLHDL
Mm-Nrf3_NP_035033 VSLIRDIRRRGKNKVAAQNCRKRKLDIILNLEDDICNLQAKKEALKNEQTQCSKAIDIMRQKLHGL
Xl-Nrf3_XP_018122887 VTVIRDIRRRGKNKVAAQNCRKRKLDVILNLDDDVCQLKARKEKLLKERAQCSKSTSHMKQKLNDL
Xl-Nrf3L_XP_018124724 VTVIRDIRRRGKNKVAAQNCRKRKLDVILNLEDDVGQLQARKEKLLKERAQCSKSMSHMKQKLNDL
Ac-Nrf3_XP_003222201 LSLIRDIRRRGKNKVAAQNCRKRKLDVIMNLKDDVCHLQTEKEKLKRQKTKRNRSISNIKQKINFL
Dr-Nrf2b_NP_001244112 HTLIRDIRRRGKNKMAAQSCRKRKMDSLFGLEDEIEDLKRKKDQCMEEKERNARELCETKEKVRKL
*Dr-Nrf1a_JX867114 HANQNTFAVEAKTSMAAQNCRRRKLDVLVELEHGVDSLRRLRAKILREKSEISCSIKEVKQRLNSL
Dr-Nrf3_NP_998396 VTLLRDVRRRGKNKLAAQNCRKRKLDAIVGLQGEVEGLMVQRDALLRQRDHTAKALSATAERFEAL
*Dm-CncC_NP_732833 LSLIRDIRRRGKNKVAAQNCRKRKLDQILTLEDEVNAVVKRKTQLNQDRDHLESERKRISNKFAML
Nv-Nach6_AOP32002 AKYVRDVRRRGKNKEAARICRKRKMDAIETLDDEITRLKQQRQSMFDERKDLQQETAELKRKISEL
Ta-Nach3_XP_002116568 AALARDVRRRGKNKFAARNCRKRKIDDIDGLKDEVDELEVQKESLLAEVKKLEEESEKYRKKSEAM
*Sp-Nach8_NP_001123278 MQLVRDIRRRGKNKVAAQNCRKRKIDAIQIVESTVGELRMERDKLVKERDSIDKEVNEMQQRYAEL
*Ob-Nach7_XP_014784776 LRLIRDIRRRGKNKVAAQNCRKRKLDIIGTLEEEVIQMRQERDRLLREGSLIHKQTREMKEKYNVR
Aq-Nach5_XP_003385261 KEEVKAIRKRGKNKNAAKNCRYRKLEHLSGLQQEIDSLKIKKSKLALKALSLQREIDGYKSKCTQK
Aq-Nach4_XP_011402550 VAFMKEWRRRGKNKMAAQIARKRKREELFELEDDMDSLRQKKSKLQQSVAKLNALIASYKRRVEVG
Es-Nach2_WP_062268396 VAVAKEWRRRNKNKSAAQVARKRKRDEAQSLQEEVDGLKKEKKELEEKLELLPGEIKQWKTKAQAL
*En-Nach1_WP_034835593 VAAMKKWRRNSKNRFSAQACRERKSKGIESLNVTIKKLKKEKKELEQKLRSLPDVIKEWKTKAQAL
*Ce-Skn1_NP_741404 RQLIRKIRRRGKNKVAARTCRQRRTDRHDKMSHYI...............................

**fig. S5**

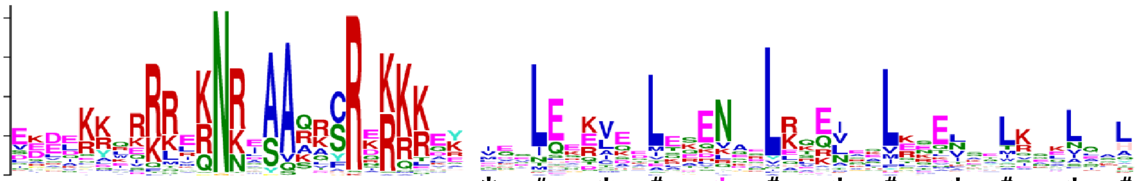

**Human vs Unclassified bZIP**

* # * # * # * # * # * #

| Group | Mark | Sequence | Alignment |
|---|---|---|---|
| CB (BATF,CHOP) | | Sc-Yap5_NP_012283 | ENDEELQKKKRQNRDAQRAYRERKNNK.LQVLEETIESLSKVVKNYETKLNRLQNELQAKESENHAL |
| | | Sc-Yap7_KZV07854 | NGVDSVEKRRRQNRDAQRAYRERRTTR.IQVLEEKVEMLHNLVDDWQRKYKLLESEFSDTKENLQKS |
| | | Sc-Cin5_NP_014671 | GKPLRNTKRAAQNRSAQKAFRQRREKY.IKNLEEKSKLFDG....LMKENSELKKMIESLKSKLKE. |
| | | Sc-Yap6_EGA87413 | GKTLRNTRRAAQNRTAQKAFRQRKEKY.IKNLEQKSKIFDD....LLAENNNFKSLNDSLRNDNNIL |
| | | Sc-Yap1_NP_013707 | LDPETKQKRTAQNRAAQRAFRERKERK.MKELEKKVQSLESIQQQNEVEATFLRDQLITLVNELKKY |
| | | Sc-Yap3_NP_011854 | VPDDSKAKKKAQNRAAQKAFRERKEAR.MKELQDKLLESERNRQSLLKEIEELRKANTEINAENRLL |
| | * (black) | Hs-CHOP_P35638 | EEDQGRTRKRKQSGHSPARAGKQRMKEKEQENERKVAQLAEENERLKQEIERLTREVEATRRALIDR |
| | | Dd-HY5L1_XP_641680 | ERHQKRQRRLVKNREAAQLFRQRQKAY.IQDLEKKVSDLTGTNSEFRARVELLNSENKLIREQLLYL |
| | | Sc-Met28p_EWG90397 | ELDKIKQERRRKNTEASQRFRIRKKQKNFENMN.KLQNLNTQINKLRDRIEQLNKENEFWKAKLSDI |
| | * (black) | Hs-BATF_AAC50314 | SDDVRRVQRREKNRIAAQKSRQRQTQK.ADTLHLESEDLEKQNAALRKEIKQLTEELKYFTSVLNSH |
| ATF (ATF2,ATF3,ATF4) | * (red) | Gh2-MEQ_YP_001033993 | EEKQKLERRRKRNRDAARRRRRKQTDY.VDKLHEACEELQRANEHLRKEIRDLRTECTSLRVQLACH |
| | * (black) | Hs-ATF2_CAA33886 | DPDEKRRKFLERNRAAASRCRQKRKVW.VQSLEKKAEDLSSLNGQLQSEVTLLRNEVAQLKQLLLAH |
| | * (black) | Hs-ATF3_AAA20506 | EEDERKKRRRERNKIAAAKCRNKKKEK.TECLQKESEKLESVNAELKAQIEELKNEKQHLIYMLNLH |
| | * (black) | Hs-ATF4_CAG30270 | EKLDKKLKKMEQNKTAATRYRQKKRAE.QEALTGECKELEKKNEALKERADSLAKEIQYLKDLIEEV |
| | * (red) | Mb-bZIP-TF1_XP_001743420 | DDDDHAGSTSNPNKSAADRYRKKKREE.FERLQHDTEAMKAENLELKTRLSKLRNEAEFLANMLQSA |
| | * (red) | Ce-bZIP-TF2_NP_500318 | DETKLLSRKRQQNKVAAARYRDKQKAK.WQDLLDQLEAEEDRNQRLKLQAGHLEKEVAEMRQAFLAK |
| | | Co-ATF2L2_XP_004364450 | DDESNKNKLRERNKTAAAKSRLKKKQR.EHQLQQRARHMVERNGDLKTAMAKMETEVQALREKMAAA |
| | | Co-ATF4L1_XP_004343869 | LDEEEKIEKRRRNKIAAARCRDKKREK.QSILDERTERMREENINLKQKVAQLEMEVSYLKNLVLAA |
| | | Ob-ATF4L1_KOF68678 | DEEIKKEKRREQNRRAAARCRNKKKWE.ERLSQANLFEEEKKQIMLKSIVRKLTDMKQNMEKLLSNV |
| CPE (C/EBP,PAR,E4BP4) | * (black) | Hs-CEBPa_CAA72289 | KNSNEYRVRRERNNIAVRKSRDKAKQR.NVETQQKVLELTSDNDRLRKRVEQLSRELDTLRGIFRQL |
| | * (red) | Mb-bZIP-TF5_XP_001743794 | EDSNNYRIKRIRNNEAVRRCRIKKKQE.MEEKAMRLELLEHKVSDLENCNRKLSELIVEQQKEIQRL |
| | | Sc-GCN4_NP_010907 | ESSDPAALKRARNTEAARRSRARKLQR.MKQLEDKVEELLSKNYHLENEVARLKKLVGER....... |
| | * (red) | Co-bZIP-TF1_XP_004363551 | ETLADYLDKRKKNNDAVKKCRARKRMA.VVATEEECQRLSGENASLRDRVGSLEAEVAYLKNLLISA |
| | * (blue) | Co-ATF4L2_XP_004349044 | DLPQRKLSRRERNNIAVRRCRDKNREK.SLAAKSQCETVAQENANLRVRIHSLEQEVSYLKSMLLSQ |
| | * (blue) | Aq-ATF7L_XP_011402548 | SKLAKKAERKEKNNAASKVSRAKRKQK.MKSLFEREKELESENARLKLQVEEMTKEAEKLKKQLILR |
| | * (red) | Ce-bZIP-TF1_NP_502961 | QKDEAYLDRRRRNNEAARKSRESRKKV.DQDNSVRVTYLERENQCLRVYVQQLQLQNESMRQHLLLQ |
| | * (black) | Hs-DBP_AAA81374 | QKDEKYWSRRYKNNEAAKRSRDARRLK.ENQISVRAAFLEKENALLRQEVVAVRQELSHYRAVLSRY |
| | | Sp-HLFL1-BRLZ1_XP_003724320 | TKDARYWVKRIKNNLSAKRSREKRRMA.DNVMESKVSKLAQENEDLRSELANIKRLVQDNIIKDTKQ |
| | | Ob-HP8-BRLZ1_XP_014769996 | RKDQCYWEKRRKNNEAARRSREKRRLH.DMALEKRIVELSRESCILRTQLYAVKKRYGIPREEPIIL |
| | | Hr-HP8L_XP_009021353 | KKDKEYWMKRQKNNAAAKKSREKRRLN.DVVLTNQIVQLTNENKRLKVELQAIKQRFGLSISSPY.. |
| | * (black) | Hs-E4BP4_AAA93067 | KKDAMYWEKRRKNNEAAKRSREKRRLN.DLVLENKLIALGEENATLKAELLSLKLKFGLISSTAYAQ |
| Fos | | Hs-cFos_CAA24756 | EEEEKRRIRRERNKMAAAKCRNRRREL.TDTLQAETDQLEDEKSALQTEIANLLKEKEKLEFILAAH |
| | * (red) | Nv-bZIP-TF1_XP_001628839 | EEEEKRRLRRERNKQAANRCRKRKRDK.IEMLERTAQEIDDSNKALETDIANMRTELTELMSVLRSH |
| | * (red) | Co-bZIP-TF3_XP_004349462 | GSIDKRSLKRLRNREAAARCRNRRRQL.IDELSTQVAELVAEKTTMAATIARLEAELATTRGN.... |
| | * (blue) | Sc-ATF2L2_Sko1p_AJT01605 | EQERKRKEFLERNRVAASKFRKRKKEY.IKKIENDLQFYESEYDDLTQVIGKLCGIIPSSSSNSQFN |
| | | Sc-HAC1_NP_116622 | EKEQRRIERILRNRRAAHQSREKKRLH.LQYLERKCSLLENLLNSVNLEKLADHEDALTCSHDAFVA |
| | | Sp-cFos_XP_003726998 | EEEVRRRLQKERNRDAASKCRSKKRNA.VGHLVEEAQQLETENMKLREEMKALESERSQLQFLLDMH |
| CMC (CNC,Maf,CREB(ATF1)) | * (black) | Hs-ATF1_AAH29619 | DPQLKREIRLMKNREAARECRRKKKEY.VKCLENRVAVLENQNKTLIEELKTLKDLYSNKSV..... |
| | * (blue) | Hr-ATF4L1_XP_009024663 | HDSLRKNRRREQNRIAARKCREKKRVQ.VDSILKGYADTLKENKKLKQETQVLKLVVNNLQNVLTSH |
| | * (red) | Co-bZIP-TF6_KJE89372 | NDTDPNQRRREKNREAAQACRIKKKVY.VNSMQGSVDSVAETNNHLNLQLSMVQQNTLRLHGFSQQL |
| | * (blue) | Co-XBP1L_XP_004347974 | ISDLKMQRRRVKNREAAQVCRKKKKSF.VVDLEGNMSVLQREQDNLRENLRTAEATFQQAKTATATK |
| | * (black) | Hs-Nrf1_NM_003204.2 | LSLIRDIRRRGKNKMAAQNCRKRKLDT.ILNLERDVEDLQRDKARLLREKVEFLRSLRQMKQKVQSL |
| | * (black) | Hs-Bach1_NP_001177 | LDCIHDIRRRSKNRIAAQRCRKRKLDC.IQNLESEIEKLQSEKESLLKERDHILSTLGETKQNLTGL |
| | | Es-Nach2_WP_062268396 | VAVAKEWRRRNKNKSAAQVARKRKRDE.AQSLQEEVDGLKKEKKELEEKLELLPGEIKQWKTKAQAL |
| | | En-Nach1_WP_034835593 | VAAMKKWRRNSKNRFSAQACRERKSKG.IESLNVTIKKLKKEKKELEQKLRSLPDVIKEWKTKAQAL |
| | * (red) | Co-bZIP-TF5_XP_004343898 | IRDLKDLRRKMKNRNAAARCRQRKRKE.TGVIRDRMSGLHVEVAFLRQENAILKSFLRTAGISLPDS |
| | * (black) | Hs-cMaf_AAC27037 | VIRLKQKRRTLKNRGYAQSCRFKRVQQ.RHVLESEKNQLLQQVDHLKQEISRLVRERDAYKEKYEKL |
| | * (red) | Hr-bZIP-TF1_XP_009009099 | VQELKKKRRQLKNRNYAKTCRHKKITK.NVSLEEEVKMLRQERIHYVNEISKYKEEIKLLKMKLEIT |
| | * (red) | Mb-bZIP-TF3_XP_001744453 | VADVKAKRRRLKNRLSARLCSNKKREK.CSELEDTNRDLLAKLRQVAQENKTLKSETNRLKEANTAL |
| JOAX (JUN,OASIS,ATF6,XBP1) | * (red) | Vb-bZIP-TF2_CEM00542 | SHLDKKEQQRLRNRISAQQSRDRKKKE.FEGLSLQVDTLTTENADLRRENVALRAENTAIAAHRDQL |
| | * (red) | Co-bZIP-TF2_XP_004347848 | PEMDKKLQRLIKNREAASQSRKRKKDQ.FDTLERDLNTIKTHNAALRSQVVALEQENAVLKADNERL |
| | * (red) | Vb-bZIP-TF3_CEM20300 | EEEERMSQQKLRNRLSAQAHRDRQKRL.MRDLQERVERLSAENQHLHRENSELKQDNARLIRDVAQL |
| | * (black) | Hs-XBP1U_NP_005071 | SPEEKALRRKLKNRVAAQTARDRKKAR.MSELEQQVVDLEEENQKLLLENQLLREKTHGLVVENQEL |
| | * (black) | Hs-ATF6_BAA34722 | IAVLRRQQRMIKNRESACQSRKKKKEY.MLGLEARLKAALSENEQLKKENGTLKRQLDEVVSENQRL |
| | * (red) | Vb-bZIP-TF1_CEL94591 | VDEIKKQRKRDQNRASAVRSRAKKKEY.YTSLEHEVEALRHEATSLRAENQLLKQQLSFLQSLVQPN |
| | * (red) | Dd-bZIP-TF1_XP_642532 | EEAKKKKIRQMQNRQSAAQYRERKKEY.LEKLETIVDNLESDRNQLLQQTKQLGMLQNENYLKINQL |
| | * (red) | Mb-bZIP-TF2_XP_001742296 | EIKEKKERRMLKNRESASLSRKRKKEY.LETLEHQLHDAQQQLGRAQHQIQQLQNDNHVLREQLANY |
| | * (black) | Hs-OASIS_BAC01278 | EKALKRVRRKIKNKISAQESRRKKKEY.VECLEKKVETFTSENNELWKKVETLENANRTLLQQLQKL |
| | * (red) | Hr-bZIP-TF2_XP_009020232 | EKNLKKIRRKIKNKISAQESRRKKKEY.LESLEKKVEQITQENSGLKKKVNVLENNNRNLIAELQKL |
| | * (red) | Mb-bZIP-TF4_XP_001750907 | SRELKRMRRKVKNKLSAKDSRRRRKEY.VTQLEEENAQLRARLVTLHDQSMARQSMPATTSSSSSTT |
| | | Dd-HY5L2_XP_638790 | EKVKKRQVRLLKNRQSAALSRSRKKEY.IANLESKAQELTHSTQELHVQYNKISSTTFETKSRLEFL |
| | * (red) | Dd-bZIP-TF2_XP_644283 | EKELKKQRRLVKNREYASQSRSRRKIY.VENIETKLQKTNQDCASIKSQLNSVKEENKALKKQLYSL |
| | * (red) | Co-bZIP-TF4_XP_004363356 | SSDESKVAKLEKNRQSARDCRKRKKQY.IGNLEAKVEFLTEENARLARQLAEFLATSTKLVPSINQP |
| | | Hs-cJun_AAA59197 | QERIKAERKRMRNRIAASKCRKRKLER.IARLEEKVKTLKAQNSELASTANMLREQVAQLKQKVMNH |
| | * (red) | Ob-bZIP-TF1_XP_014790770 | FSRNLQMIRRKKNRLAAQKCREKKKER.IRILEEEIKSLIKENYSLKQANYELGEKLSEQQKKLEEA |

### fig. S6

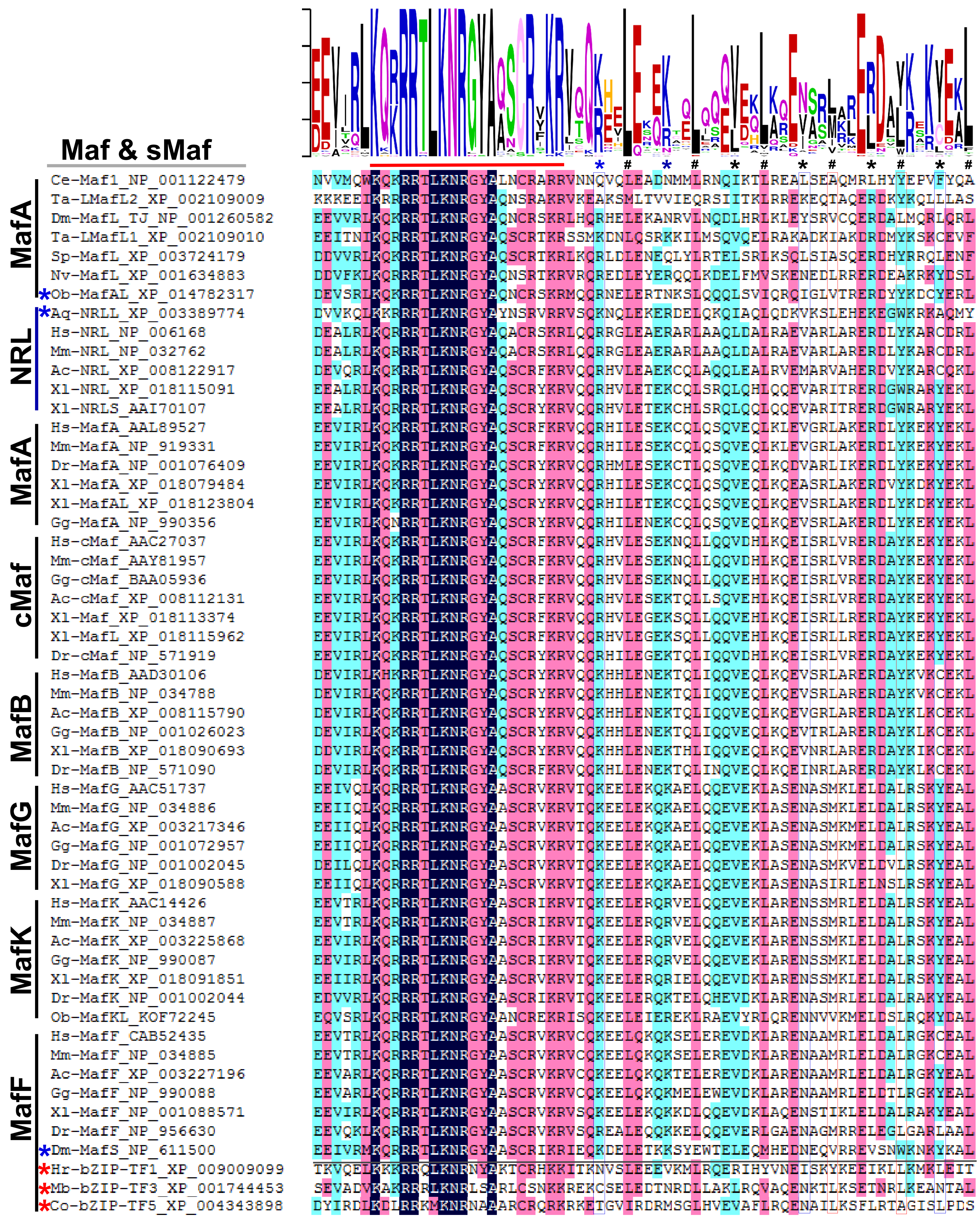
Maf & sMaf
MafA
NRL
MafA
cMaf
MafB
MafG
MafK
MafF
Ce-Maf1_NP_001122479
Ta-LMafL2_XP_002109009
Dm-MafL_TJ_NP_001260582
Ta-LMafL1_XP_002109010
Sp-MafL_XP_003724179
Nv-MafL_XP_001634883
*Ob-MafAL_XP_014782317
*Aq-NRLL_XP_003389774
Hs-NRL_NP_006168
Mm-NRL_NP_032762
Ac-NRL_XP_008122917
Xl-NRL_XP_018115091
Xl-NRLS_AAI70107
Hs-MafA_AAL89527
Mm-MafA_NP_919331
Dr-MafA_NP_001076409
Xl-MafA_XP_018079484
Xl-MafAL_XP_018123804
Gg-MafA_NP_990356
Hs-cMaf_AAC27037
Mm-cMaf_AAY81957
Gg-cMaf_BAA05936
Ac-cMaf_XP_008112131
Xl-Maf_XP_018113374
Xl-MafL_XP_018115962
Dr-cMaf_NP_571919
Hs-MafB_AAD30106
Mm-MafB_NP_034788
Ac-MafB_XP_008115790
Gg-MafB_NP_001026023
Xl-MafB_XP_018090693
Dr-MafB_NP_571090
Hs-MafG_AAC51737
Mm-MafG_NP_034886
Ac-MafG_XP_003217346
Gg-MafG_NP_001072957
Dr-MafG_NP_001002045
Xl-MafG_XP_018090588
Hs-MafK_AAC14426
Mm-MafK_NP_034887
Ac-MafK_XP_003225868
Gg-MafK_NP_990087
Xl-MafK_XP_018091851
Dr-MafK_NP_001002044
Ob-MafKL_KOF72245
Hs-MafF_CAB52435
Mm-MafF_NP_034885
Ac-MafF_XP_003227196
Gg-MafF_NP_990088
Xl-MafF_NP_001088571
Dr-MafF_NP_956630
*Dm-MafS_NP_611500
*Hr-bZIP-TF1_XP_009009099
*Mb-bZIP-TF3_XP_001744453
*Co-bZIP-TF5_XP_004343898

fig. S7

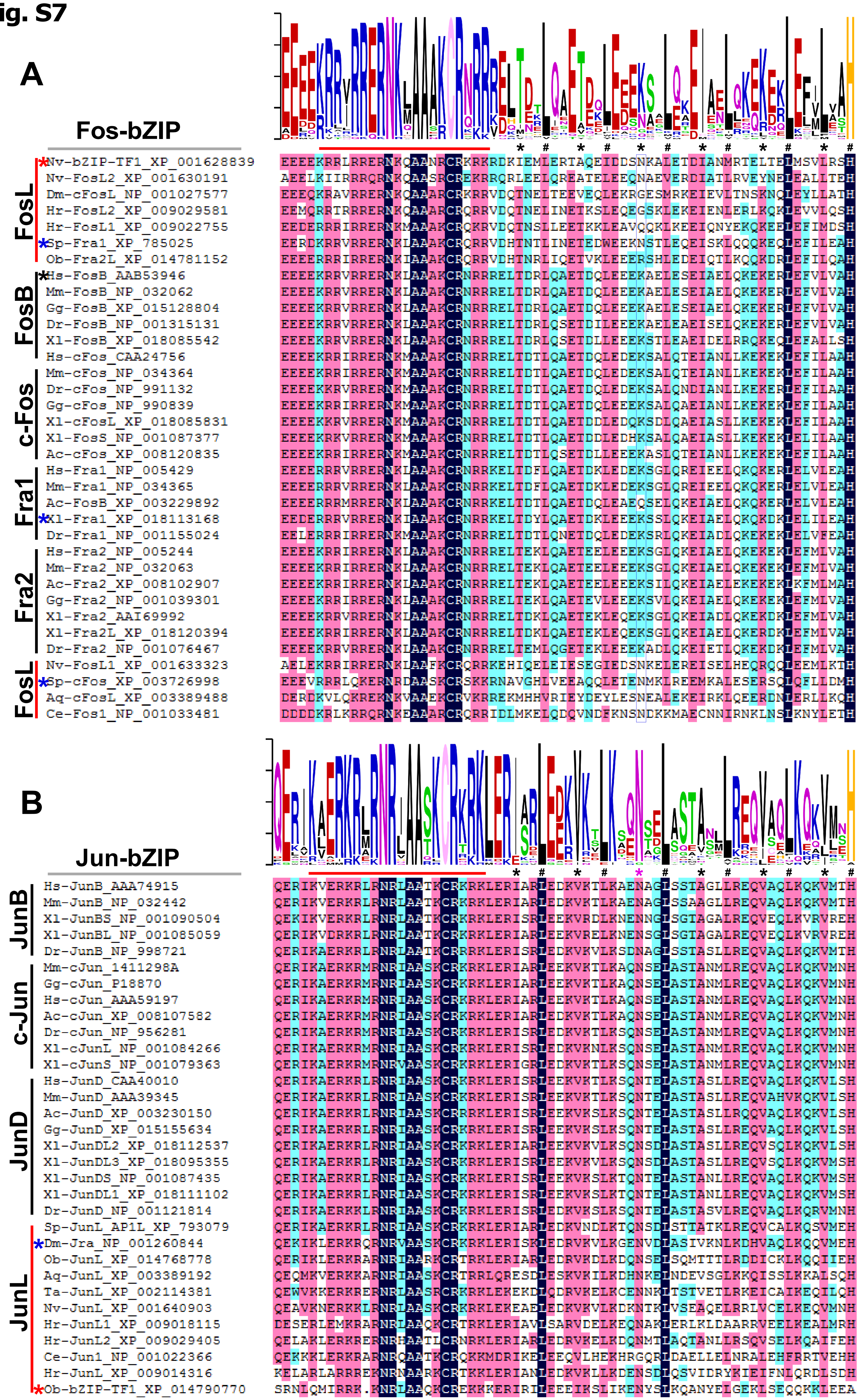
A
Fos-bZIP
FosL
Nv-bZIP-TF1_XP_001628839
Nv-FosL2_XP_001630191
Dm-cFosL_NP_001027577
Hr-FosL2_XP_009029581
Hr-FosL1_XP_009022755
Sp-Fra1_XP_785025
Ob-Fra2L_XP_014781152
FosB
Hs-FosB_AAB53946
Mm-FosB_NP_032062
Gg-FosB_XP_015128804
Dr-FosB_NP_001315131
Xl-FosB_XP_018085542
c-Fos
Hs-cFos_CAA24756
Mm-cFos_NP_034364
Dr-cFos_NP_991132
Gg-cFos_NP_990839
Xl-cFosL_XP_018085831
Xl-FosS_NP_001087377
Ac-cFos_XP_008120835
Fra1
Hs-Fra1_NP_005429
Mm-Fra1_NP_034365
Ac-FosB_XP_003229892
Xl-Fra1_XP_018113168
Dr-Fra1_NP_001155024
Fra2
Hs-Fra2_NP_005244
Mm-Fra2_NP_032063
Ac-Fra2_XP_008102907
Gg-Fra2_NP_001039301
Xl-Fra2_AAI69992
Xl-Fra2L_XP_018120394
Dr-Fra2_NP_001076467
FosL
Nv-FosL1_XP_001633323
Sp-cFos_XP_003726998
Aq-cFosL_XP_003389488
Ce-Fos1_NP_001033481
B
Jun-bZIP
JunB
Hs-JunB_AAA74915
Mm-JunB_NP_032442
Xl-JunBS_NP_001090504
Xl-JunBL_NP_001085059
Dr-JunB_NP_998721
c-Jun
Mm-cJun_1411298A
Gg-cJun_P18870
Hs-cJun_AAA59197
Ac-cJun_XP_008107582
Dr-cJun_NP_956281
Xl-cJunL_NP_001084266
Xl-cJunS_NP_001079363
JunD
Hs-JunD_CAA40010
Mm-JunD_AAA39345
Ac-JunD_XP_003230150
Gg-JunD_XP_015155634
Xl-JunDL2_XP_018112537
Xl-JunDL3_XP_018095355
Xl-JunDS_NP_001087435
Xl-JunDL1_XP_018111102
Dr-JunD_NP_001121814
JunL
Sp-JunL_AP1L_XP_793079
Dm-Jra_NP_001260844
Ob-JunL_XP_014768778
Aq-JunL_XP_003389192
Ta-JunL_XP_002114381
Nv-JunL_XP_001640903
Hr-JunL1_XP_009018115
Hr-JunL2_XP_009029405
Ce-Jun1_NP_001022366
Hr-JunL_XP_009014316
Ob-bZIP-TF1_XP_014790770

fig. S8

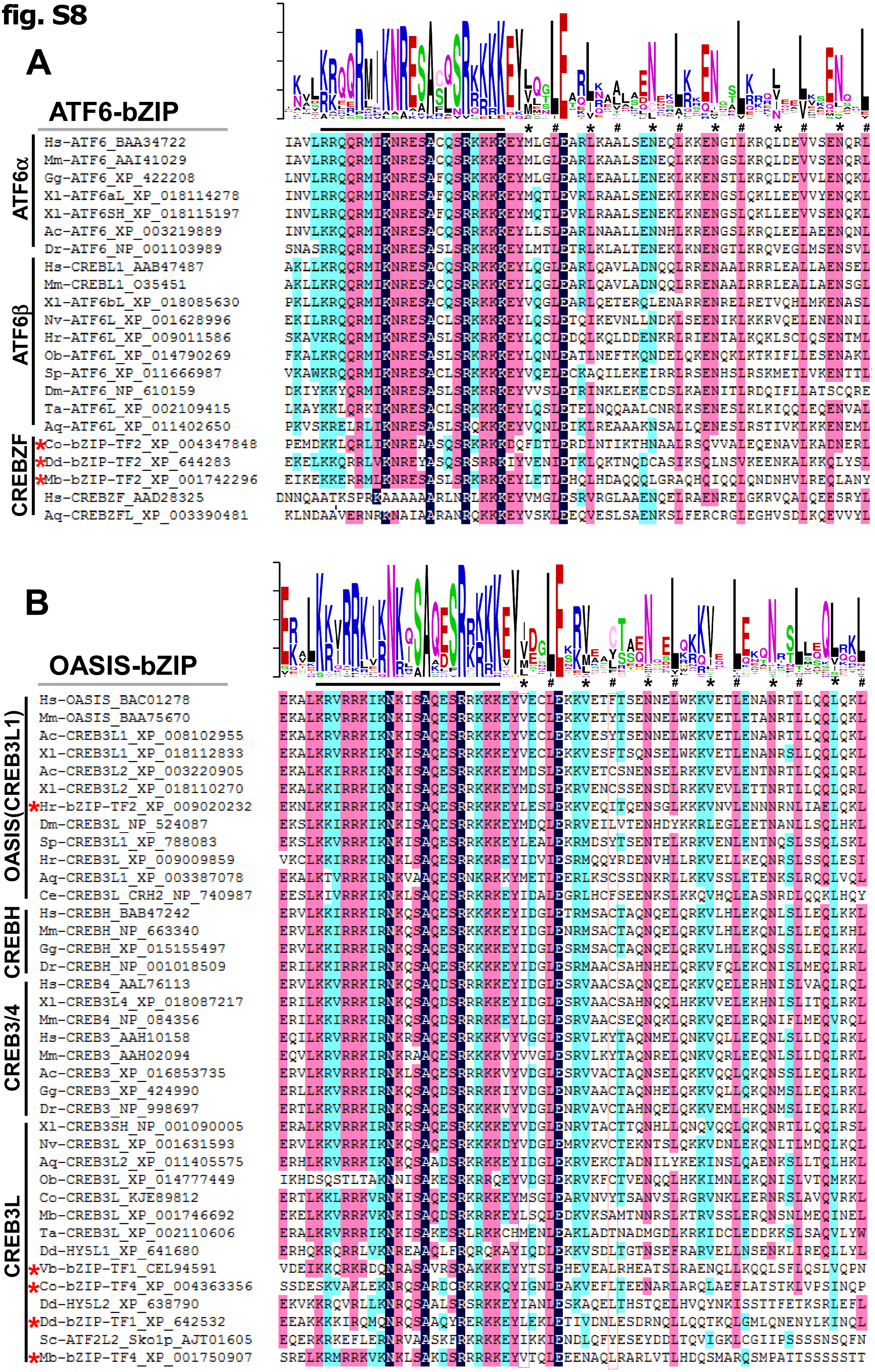
A
ATF6-bZIP
ATF6α
Hs-ATF6_BAA34722
Mm-ATF6_AAI41029
Gg-ATF6_XP_422208
Xl-ATF6aL_XP_018114278
Xl-ATF6SH_XP_018115197
Ac-ATF6_XP_003219889
Dr-ATF6_NP_001103989
ATF6β
Hs-CREBL1_AAB47487
Mm-CREBL1_O35451
Xl-ATF6bL_XP_018085630
Nv-ATF6L_XP_001628996
Hr-ATF6L_XP_009011586
Ob-ATF6L_XP_014790269
Sp-ATF6_XP_011666987
Dm-ATF6_NP_610159
Ta-ATF6L_XP_002109415
Aq-ATF6L_XP_011402650
CREBZF
*Co-bZIP-TF2_XP_004347848
*Dd-bZIP-TF2_XP_644283
*Mb-bZIP-TF2_XP_001742296
Hs-CREBZF_AAD28325
Aq-CREBZFL_XP_003390481
B
OASIS-bZIP
OASIS(CREB3L1)
Hs-OASIS_BAC01278
Mm-OASIS_BAA75670
Ac-CREB3L1_XP_008102955
Xl-CREB3L1_XP_018112833
Ac-CREB3L2_XP_003220905
Xl-CREB3L2_XP_018110270
*Hr-bZIP-TF2_XP_009020232
Dm-CREB3L_NP_524087
Sp-CREB3L1_XP_788083
Hr-CREB3L_XP_009009859
Aq-CREB3L1_XP_003387078
Ce-CREB3L_CRH2_NP_740987
CREBH
Hs-CREBH_BAB47242
Mm-CREBH_NP_663340
Gg-CREBH_XP_015155497
Dr-CREBH_NP_001018509
CREB3/4
Hs-CREB4_AAL76113
Xl-CREB3L4_XP_018087217
Mm-CREB4_NP_084356
Hs-CREB3_AAH10158
Mm-CREB3_AAH02094
Ac-CREB3_XP_016853735
Gg-CREB3_XP_424990
Dr-CREB3_NP_998697
CREB3L
Xl-CREB3SH_NP_001090005
Nv-CREB3L_XP_001631593
Aq-CREB3L2_XP_011405575
Ob-CREB3L_XP_014777449
Co-CREB3L_KJE89812
Mb-CREB3L_XP_001746692
Ta-CREB3L_XP_002110606
Dd-HY5L1_XP_641680
*Vb-bZIP-TF1_CEL94591
*Co-bZIP-TF4_XP_004363356
Dd-HY5L2_XP_638790
*Dd-bZIP-TF1_XP_642532
Sc-ATF2L2_Sko1p_AJT01605
*Mb-bZIP-TF4_XP_001750907

fig. S9

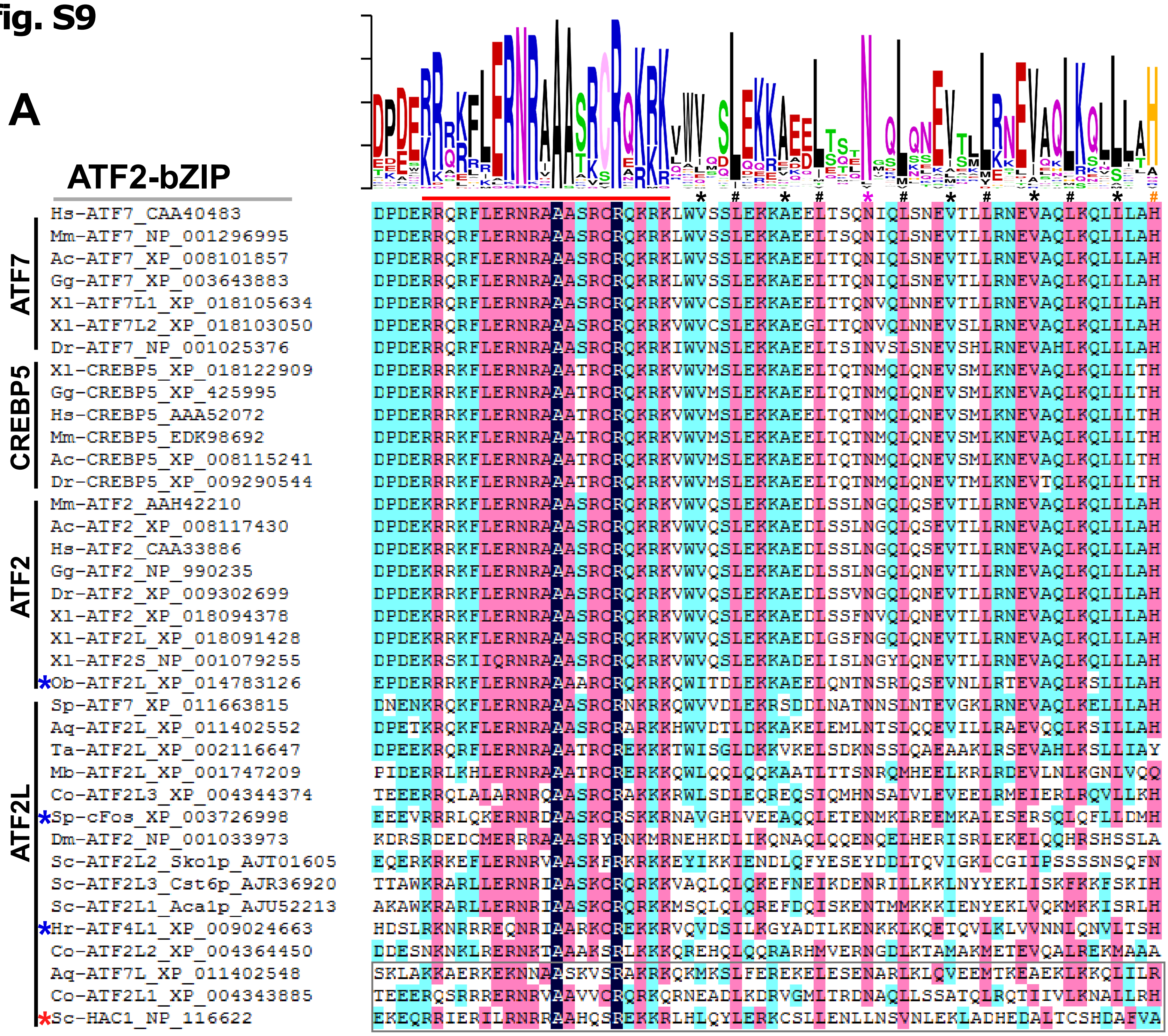
A
ATF2-bZIP
ATF7
CREBP5
ATF2
ATF2L
Hs-ATF7_CAA40483
Mm-ATF7_NP_001296995
Ac-ATF7_XP_008101857
Gg-ATF7_XP_003643883
Xl-ATF7L1_XP_018105634
Xl-ATF7L2_XP_018103050
Dr-ATF7_NP_001025376
Xl-CREBP5_XP_018122909
Gg-CREBP5_XP_425995
Hs-CREBP5_AAA52072
Mm-CREBP5_EDK98692
Ac-CREBP5_XP_008115241
Dr-CREBP5_XP_009290544
Mm-ATF2_AAH42210
Ac-ATF2_XP_008117430
Hs-ATF2_CAA33886
Gg-ATF2_NP_990235
Dr-ATF2_XP_009302699
Xl-ATF2_XP_018094378
Xl-ATF2L_XP_018091428
Xl-ATF2S_NP_001079255
*Ob-ATF2L_XP_014783126
Sp-ATF7_XP_011663815
Aq-ATF2L_XP_011402552
Ta-ATF2L_XP_002116647
Mb-ATF2L_XP_001747209
Co-ATF2L3_XP_004344374
*Sp-cFos_XP_003726998
Dm-ATF2_NP_001033973
Sc-ATF2L2_Sko1p_AJT01605
Sc-ATF2L3_Cst6p_AJR36920
Sc-ATF2L1_Aca1p_AJU52213
*Hr-ATF4L1_XP_009024663
Co-ATF2L2_XP_004364450
Aq-ATF7L_XP_011402548
Co-ATF2L1_XP_004343885
*Sc-HAC1_NP_116622

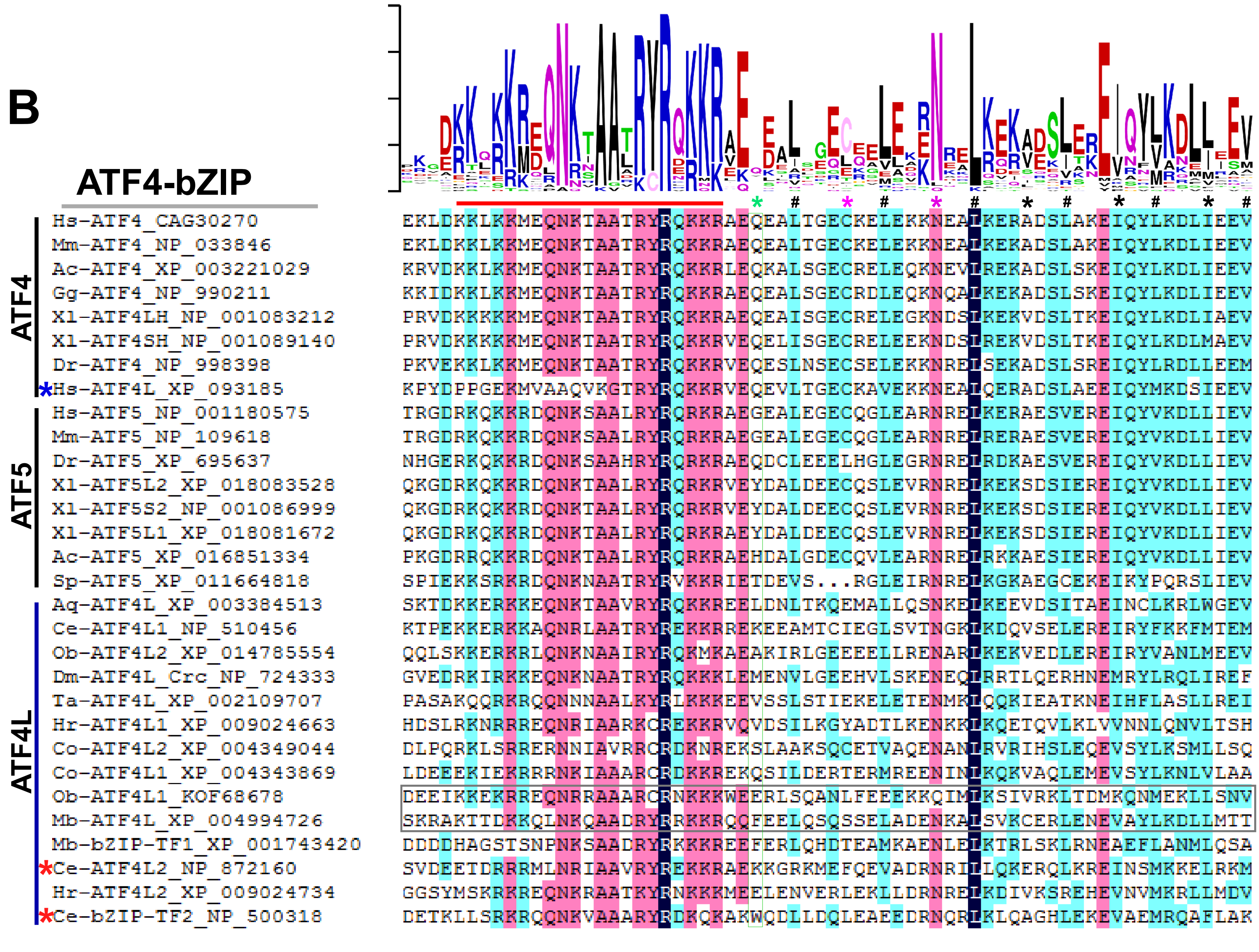
B
ATF4-bZIP
ATF4
ATF5
ATF4L
Hs-ATF4_CAG30270
Mm-ATF4_NP_033846
Ac-ATF4_XP_003221029
Gg-ATF4_NP_990211
Xl-ATF4LH_NP_001083212
Xl-ATF4SH_NP_001089140
Dr-ATF4_NP_998398
*Hs-ATF4L_XP_093185
Hs-ATF5_NP_001180575
Mm-ATF5_NP_109618
Dr-ATF5_XP_695637
Xl-ATF5L2_XP_018083528
Xl-ATF5S2_NP_001086999
Xl-ATF5L1_XP_018081672
Ac-ATF5_XP_016851334
Sp-ATF5_XP_011664818
Aq-ATF4L_XP_003384513
Ce-ATF4L1_NP_510456
Ob-ATF4L2_XP_014785554
Dm-ATF4L_Crc_NP_724333
Ta-ATF4L_XP_002109707
Hr-ATF4L1_XP_009024663
Co-ATF4L2_XP_004349044
Co-ATF4L1_XP_004343869
Ob-ATF4L1_KOF68678
Mb-ATF4L_XP_004994726
Mb-bZIP-TF1_XP_001743420
*Ce-ATF4L2_NP_872160
Hr-ATF4L2_XP_009024734
*Ce-bZIP-TF2_NP_500318

fig. S10

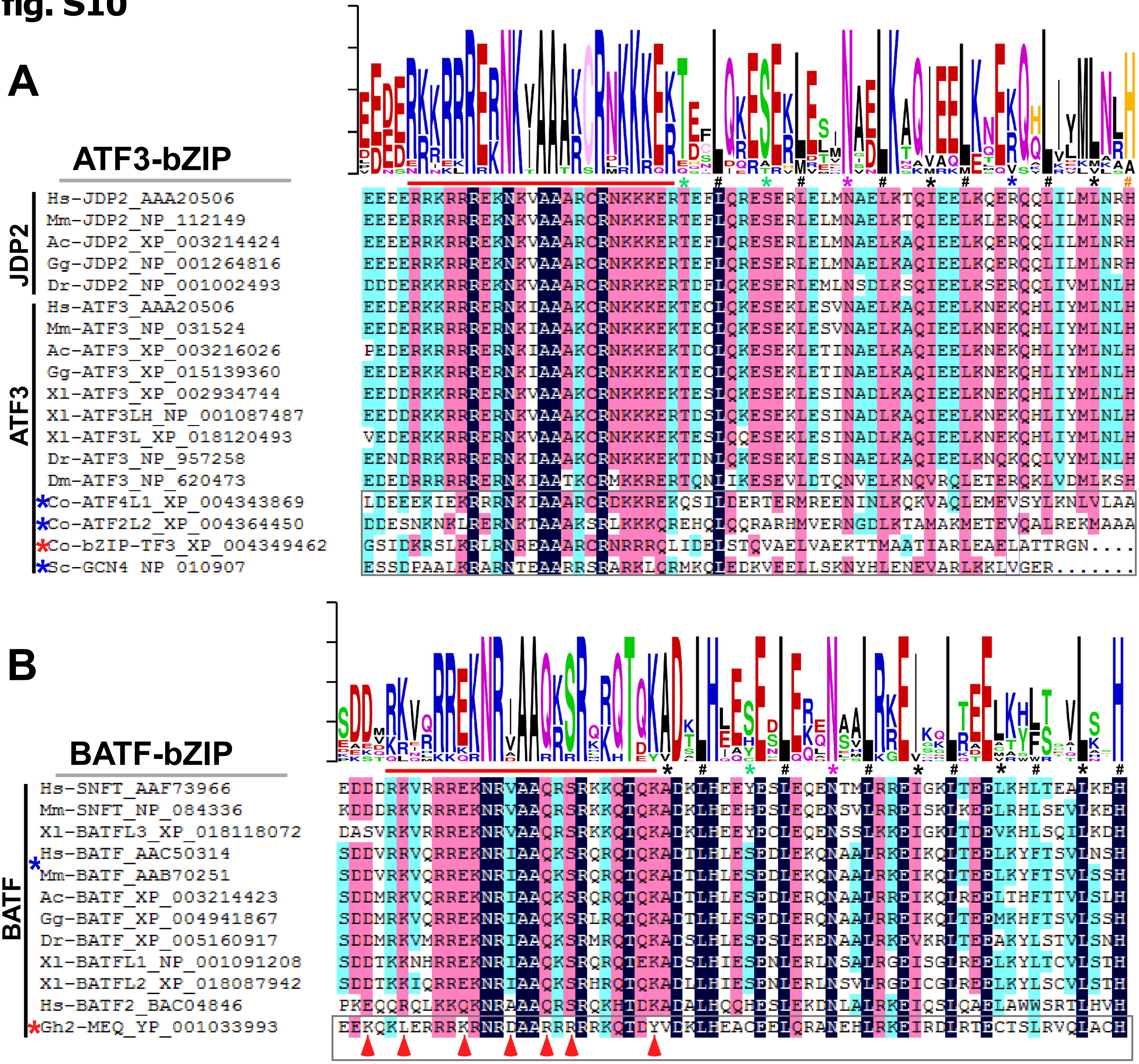

A
ATF3-bZIP
JDP2
ATF3
Hs-JDP2_AAA20506
Mm-JDP2_NP_112149
Ac-JDP2_XP_003214424
Gg-JDP2_NP_001264816
Dr-JDP2_NP_001002493
Hs-ATF3_AAA20506
Mm-ATF3_NP_031524
Ac-ATF3_XP_003216026
Gg-ATF3_XP_015139360
Xl-ATF3_XP_002934744
Xl-ATF3LH_NP_001087487
Xl-ATF3L_XP_018120493
Dr-ATF3_NP_957258
Dm-ATF3_NP_620473
Co-ATF4L1_XP_004343869
Co-ATF2L2_XP_004364450
Co-bZIP-TF3_XP_004349462
Sc-GCN4_NP_010907
B
BATF-bZIP
BATF
Hs-SNFT_AAF73966
Mm-SNFT_NP_084336
Xl-BATFL3_XP_018118072
Hs-BATF_AAC50314
Mm-BATF_AAB70251
Ac-BATF_XP_003214423
Gg-BATF_XP_004941867
Dr-BATF_XP_005160917
Xl-BATFL1_NP_001091208
Xl-BATFL2_XP_018087942
Hs-BATF2_BAC04846
Gh2-MEQ_YP_001033993

fig. S11

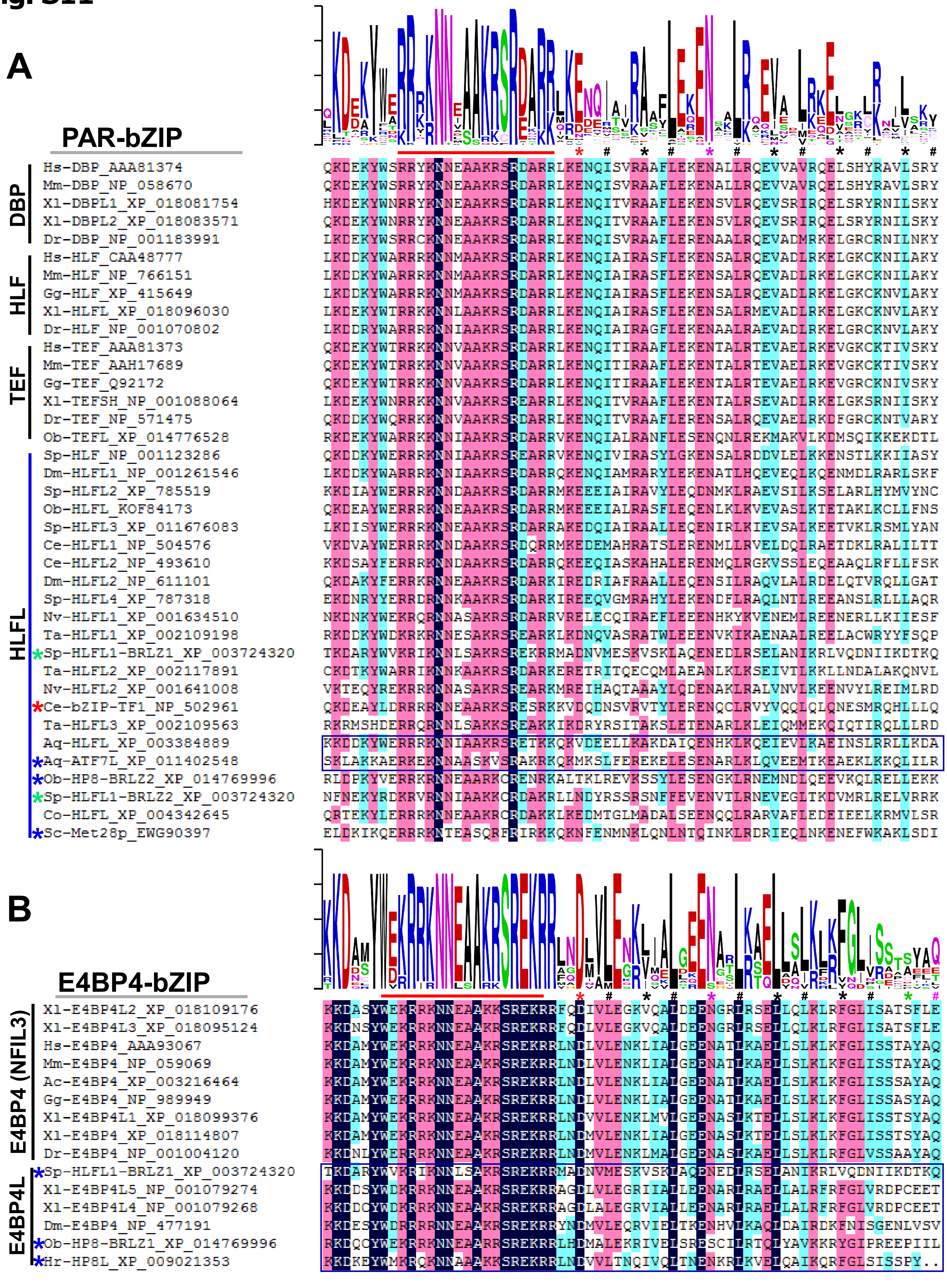
A
PAR-bZIP
DBP
HLF
TEF
HLFL
Hs-DBP_AAA81374 QKDEKYWSRRYKNNEAAKRSRDARRLKENQISVRAAFLEKENALLRQEVVAVRQELSHYRAVLSRY
Mm-DBP_NP_058670 QKDEKYWSRRYKNNEAAKRSRDARRLKENQISVRAAFLEKENALLRQEVVAVRQELSHYRAVLSRY
Xl-DBPL1_XP_018081754 HKDEKYWNRRYKNNEAAKRSRDARRLKENQITVRAAFLEKENSVLRQEVSRIRQELSRYRNILSKY
Xl-DBPL2_XP_018083571 QKDEKYWNRRYKNNEAAKRSRDARRLKENQITVRAAFLEKENSVLRQEVSRIRQELSRYRNILSKY
Dr-DBP_NP_001183991 LKDEKYWSRRCKNNEAAKRSRDARRLKENQISVRAAFLERENAALRQEVADMRKELGRCRNILNKY
Hs-HLF_CAA48777 LKDDKYWARRRKNNMAAKRSRDARRLKENQIAIRASFLEKENSALRQEVADLRKELGKCKNILAKY
Mm-HLF_NP_766151 LKDDKYWARRRKNNMAAKRSRDARRLKENQIAIRASFLEKENSALRQEVADLRKELGKCKNILAKY
Gg-HLF_XP_415649 LKDDKYWARRRKNNMAAKRSRDARRLKENQIAIRASFLEKENSALRQEVADLRKELGKCKNVLAKY
Xl-HLFL_XP_018096030 LKDEKYWTRRKKNNLAAKRSRDARRLKENQIAIRASFLEKENSALRMEVADLRKELGKCKNVLAKY
Dr-HLF_NP_001070802 LKDDRYWARRRKNNIAAKRSRDARRLKENQIAIRAGFLEKENAALRAEVADLRKELGRCKNVLAKY
Hs-TEF_AAA81373 QKDEKYWTRRKKNNVAAKRSRDARRLKENQITIRAAFLEKENTALRTEVAELRKEVGKCKTIVSKY
Mm-TEF_AAH17689 QKDEKYWTRRKKNNVAAKRSRDARRLKENQITIRAAFLEKENTALRTEVAELRKEVGKCKTIVSKY
Gg-TEF_Q92172 QKDEKYWTRRKKNNVAAKRSRDARRLKENQITIRAAFLEKENTALRTEVAELRKEVGRCKNIVSKY
Xl-TEFSH_NP_001088064 LKDEKYWNRRKKNNVAAKRSREARRLKENQITVRAAFLEKENTALRSEVADLRKELGKSRNIISKY
Dr-TEF_NP_571475 QKDDKYWQRRKKNNVAAKRSRDARRLKENQITVRAAFLERENSALRQEVAELRKDFGRCKNTVARY
Ob-TEFL_XP_014776528 RKDEKYWARRKKNNIAAKRSRDARRVKENQIALRANFLESENQNLREKMAKVLKDMSQIKKEKDTL
Sp-HLF_NP_001123286 QKDDKYWERRRKNNIAAKRSREARRVKENQIVIRASYLGKENSALRDDVLELKKENSTLKKIIASY
Dm-HLFL1_NP_001261546 LKDDKYWARRRKNNIAAKRSRDARRQKENQIAMRARYLEKENATLHQEVEQLKQENMDLRARLSKF
Sp-HLFL2_XP_785519 KKDIAYWERRRKNNDAAKRSRDARRMKEEEIAIRAVYLEQDNMKLRAEVSILKSELARLHYMVYNC
Ob-HLFL_KOF84173 QKDEAYWERRRKNNEAAKRSRDARRMKEEEIALRASFLEQENLKLKVEVASLKTETAKLKCLLFNS
Sp-HLFL3_XP_011676083 LKDISYWERRRKNNEAAKRSRDARRAKEDQIAIRAALLEQENIRLKIEVSALKEETVKLRSMLYAN
Ce-HLFL1_NP_504576 VKDVAYWERRRKNNDAAKRSRDQRRMKEDEMAHRATSLERENMLLRVELDQLRAETDKLRALILTT
Ce-HLFL2_NP_493610 KKDSAYFERRRKNNDAAKRSRDARRQKEEQIASKAHALERENMQLRGKVSSLEQEAAQLRFLLFSK
Dm-HLFL2_NP_611101 QKDAKYFERRKRNNEAAKRSRDARKIREDRIAFRAALLEQENSILRAQVLALRDELQTVRQLLGAT
Sp-HLFL4_XP_787318 EKDNRYYERRDRNNKAAKRSRDARKIREEQVGMRAHYLEKENDFLRAQLNTLREEANSLRLLAQR
Nv-HLFL1_XP_001634510 NKDNKYWEKRQRNNASAKRSRDARRVRELECQIRAEFLEEENHKYKVENEMLREENERLLKIIESF
Ta-HLFL1_XP_002109198 RKDDKYWDKRKRNNESAKRSREARKLKDNQVASRATWLEEENVKIKAENAALREELACWRYYFSQP
*Sp-HLFL1-BRLZ1_XP_003724320 TKDARYWVKRIKNNLSAKRSREKRRMADNVMESKVSKLAQENEDLRSELANIKRLVQDNIIKDTKQ
Ta-HLFL2_XP_002117891 CKDTKYWARRIKNNKAAKRSRDARKERETRITQECQMLAEANLKLKSEIVTLKKLLNDALAKQNVL
Nv-HLFL2_XP_001641008 VKTEQYREKRRKNNASAKRSREARKMREIHAQTAAYLQDENAKLRALVNVLKEENVYLREIMLRD
*Ce-bZIP-TF1_NP_502961 QKDEAYLDRRRRNNEAARKSRESRKKVDQDNSVRVTYLERENQCLRVYVQQLQLQNESMRQHLLLQ
Ta-HLFL3_XP_002109563 RKRMSHDERRQRNNLSAKKSREAKKIKDRYRSITAKSLETENARLKLEIQMMEKQIQTIRQLLRD
Aq-HLFL_XP_003384889 KKDDKYWERRRKNNIAAKRSRETKKQKVDEELLKAKDAIQENHKLKQEIEVLKAEINSLRRLLKDA
*Aq-ATF7L_XP_011402548 SKLAKKAERKEKNNAASKVSRAKRKQKMKSLFEREKELESENARLKLQVEEMTKEAEKLKKQLILR
*Ob-HP8-BRLZ2_XP_014769996 RLDPKYVERRKRNNEAARKCRENRKALTKLREVKSSYLESENGKLRNEMNDLQEEVKQLRELLEKK
*Sp-HLFL1-BRLZ2_XP_003724320 NFNEKYRDKRVRNNIAAKKCRDAKRLLNDYRSSRSNFFEVENVTLRNEVEGLTKDVMRLRELVRRK
Co-HLFL_XP_004342645 QRTEKYLERRRKNNEAAKRCRDAKKLKEDMTGLMADALSEENQQLRARVAFLEDEIEELKRMVLSR
*Sc-Met28p_EWG90397 ELDKIKQERRRKNTEASQRFRIRKKQKNFENMNKLQNLNTQINKLRDRIEQLNKENEFWKAKLSDI
B
E4BP4-bZIP
E4BP4 (NFIL3)
E4BP4L
Xl-E4BP4L2_XP_018109176 KKDASYWEKRRKNNEAAKKSREKRRFQDIVLEGKVQALDEENGRLRSELLQLKLRFGLISATSFLE
Xl-E4BP4L3_XP_018095124 KKDNSYWEKRRKNNEAAKRSREKRRFQDIVLEGKVQALDEENGRLRSELLQLKLRFGLISATSFLE
Hs-E4BP4_AAA93067 KKDAMYWEKRRKNNEAAKRSREKRRLNDLVLENKLIALGEENATLKAELLSLKLKFGLISSTAYAQ
Mm-E4BP4_NP_059069 KKDAMYWEKRRKNNEAAKRSREKRRLNDLVLENKLIALGEENATLKAELLSLKLKFGLISSTAYAQ
Ac-E4BP4_XP_003216464 KKDAMYWEKRRKNNEAAKRSREKRRLNDLVLENKLIALGEENATLKAELLSLKLKFGLISSSAYAQ
Gg-E4BP4_NP_989949 KKDAMYWEKRRKNNEAAKRSREKRRLNDLVLENKLIALGEENATLKAELLSLKLKFGLISSASYAQ
Xl-E4BP4L1_XP_018099376 KKDAMYWEKRRKNNEAAKRSREKRRLNDVVLENKLMVLGEENASLKTELLSLKLKFGLISSTSYAQ
Xl-E4BP4_XP_018114807 KKDAMYWEKRRKNNEAAKRSREKRRLNDMVLENKLIALGEENASLKTELLSLKLKFGLISSTSYAQ
Dr-E4BP4_NP_001004120 KKDNLYWERRRKNNEAAKRSREKRRLNDMVLENKLMALGEENASLKAELLSLKLRFGLVSSAAYAQ
*Sp-HLFL1-BRLZ1_XP_003724320 TKDARYWVKRIKNNLSAKRSREKRRMADNVMESKVSKLAQENEDLRSELANIKRLVQDNIIKDTKQ
Xl-E4BP4L5_NP_001079274 KKDDSYWDKRRKNNEAAKRSREKRRAGDLVLEGRIIALLEENARLRAELLALRFRFGLVRDPCEET
Xl-E4BP4L4_NP_001079268 KKDDCYWDKRRKNNEAAKRSREKRRAGDLALEGRVIALLEENARLRAELLALRFRFGLVRDPCEET
Dm-E4BP4_NP_477191 KKDESYWDRRRRNNEAAKRSREKRRYNDMVLEQRVIELTKENHVLKAQLDAIRDKFNISGENLVSV
*Ob-HP8-BRLZ1_XP_014769996 RKDQCYWEKRRKNNEAARRSREKRRLHDMALEKRIVELSREESCILRTQLYAVKKRYGIPREEPIIL
*Hr-HP8L_XP_009021353 KKDKEYWMKRQKNNAAAKKSREKRRLNDVVLTNQIVQLTNENKRLKVELQAIKQRFGLSISSPY..

**fig. S12**

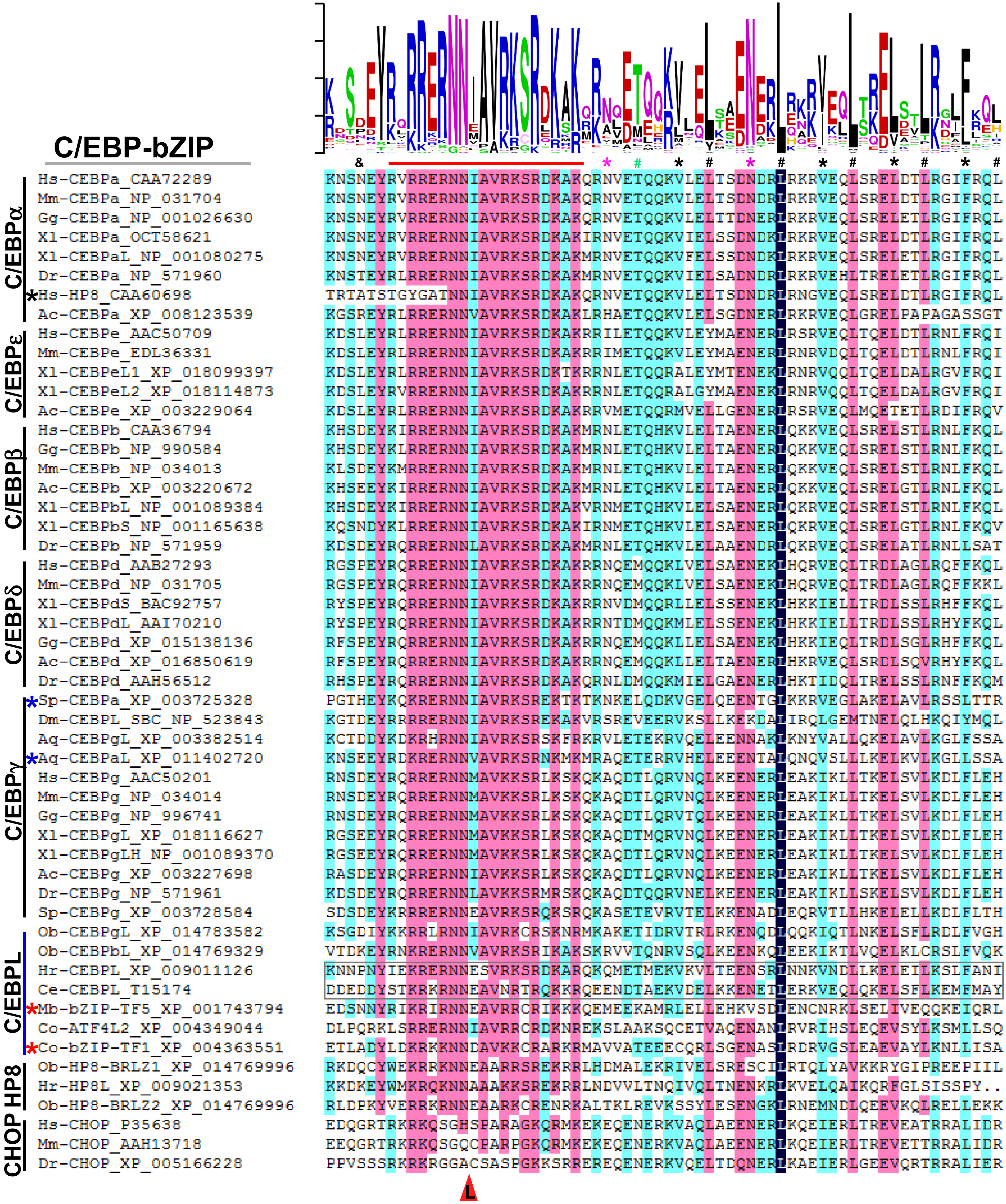
C/EBP-bZIP
C/EBPα
C/EBPε
C/EBPβ
C/EBPδ
C/EBPγ
C/EBPL
HP8
CHOP
& * # * # * # * # * # * #
Hs-CEBPa_CAA72289 KNSNEYRVRRERNNIAVRKSRDKAKQRNVETQQKVLELTSDNDRLRKRVEQLSRELDTLRGIFRQL
Mm-CEBPa_NP_031704 KNSNEYRVRRERNNIAVRKSRDKAKQRNVETQQKVLELTSDNDRLRKRVEQLSRELDTLRGIFRQL
Gg-CEBPa_NP_001026630 KNSNEYRVRRERNNIAVRKSRDKAKQRNVETQQKVLELTTDNERLRKRVEQLSRELETLRGIFRQL
Xl-CEBPa_OCT58621 KNSNEYRVRRERNNIAVRKSRDKAKIRNVETQQKVIELSSDNDKLRKRVEQLSRELDTLRGIFRQL
Xl-CEBPaL_NP_001080275 KNSNEYRVRRERNNIAVRKSRDKAKMRNVETQQKVFELSSDNDKLRKRVEQLSRELETLRGIFRQL
Dr-CEBPa_NP_571960 KNSTEYRLRRERNNIAVRKSRDKAKMRNVETQQKVIELSADNDRLRKRVEHLTRELETLRGIFRQL
*Hs-HP8_CAA60698 TRTATSTGYGATNNIAVRKSRDKAKQRNVETQQKVLELTSDNDRLRNGVEQLSRELDTLRGIFRQL
Ac-CEBPa_XP_008123539 KGSREYRLRRERNNVAVRKSRDKAKLRHAETQQKVLELSGDNERLRKRVEQLGRELPAPAGASSGT
Hs-CEBPe_AAC50709 KDSLEYRLRRERNNIAVRKSRDKAKRRILETQQKVLEYMAENERLRSRVEQLTQELDTLRNLFRQI
Mm-CEBPe_EDL36331 KDSLEYRLRRERNNIAVRKSRDKAKRRIMETQQKVLEYMAENERLRNRVDQLTQELDTLRNLFRQI
Xl-CEBPeL1_XP_018099397 KDSLEYRLRRERNNIAVRKSRDKTKRRNLETQQRALEYMTENEKLRNRVQQLTQELDALRGVFRQI
Xl-CEBPeL2_XP_018114873 KDSLEYRVRRERNNIAVRKSRDKAKRRNLETQQRALGYMAENEKLRNRVQQLTQELDALRGVFRQI
Ac-CEBPe_XP_003229064 KDSLEYRLRRERNNIAVRKSRDKAKRRVMETQQRMVELLGENERLRSRVEQLMQETETLRDIFRQV
Hs-CEBPb_CAA36794 KHSDEYKIRRERNNIAVRKSRDKAKMRNLETQHKVLELTAENERLQKKVEQLSRELSTLRNLFKQL
Gg-CEBPb_NP_990584 KHSDEYKLRRERNNIAVRKSRDKAKMRNLETQHKVLELTAENERLQKKVEQLSRELSTLRNLFKQL
Mm-CEBPb_NP_034013 KLSDEYKMRRERNNIAVRKSRDKAKMRNLETQHKVLELTAENERLQKKVEQLSRELSTLRNLFKQL
Ac-CEBPb_XP_003220672 KHSEEYKIRRERNNIAVRKSRDKAKMRNLETQHKVLELTAENERLQKKVEQLSRELGTLRNLFKQL
Xl-CEBPbL_NP_001089384 KHSDEYKIRRERNNIAVRKSRDKAKVRNMETQHKVLELSAENERLQKRVEQLSRELSTLRNLFKQL
Xl-CEBPbS_NP_001165638 KQSNDYKLRRERNNIAVRKSRDKAKIRNMETQHKVLELSAENERLQKRVEQLSRELGTLRNLFKQV
Dr-CEBPb_NP_571959 KDSDEYRQRRERNNLAVRKSRDKAKMRNLETQHKVLELAAENDRLQKRVEQLSRELATLRNLLSAT
Hs-CEBPd_AAB27293 RGSPEYRQRRERNNIAVRKSRDKAKRRNQEMQQKLVELSAENEKLHQRVEQLTRDLAGLRQFFKQL
Mm-CEBPd_NP_031705 RGSPEYRQRRERNNIAVRKSRDKAKRRNQEMQQKLVELSAENEKLHQRVEQLTRDLAGLRQFFKKL
Xl-CEBPdS_BAC92757 RYSPEYRQRRERNNIAVRKSRDKAKRRNVDMQQRLLELSSENEKLHKKIELLTRDLSSLRHFFKQL
Xl-CEBPdL_AAI70210 RYSPEYRQRRERNNIAVRKSRDKAKRRNTDMQQKMLELSSENEKLHKKIELLTRDLSSLRHYFKQL
Gg-CEBPd_XP_015138136 RFSPEYRQRRERNNIAVRKSRDKAKRRNQEMQQKLLELSAENEKLHKKIEQLTRDLSGLRHFFKQL
Ac-CEBPd_XP_016850619 RFSPEYRQRRERNNIAVRKSRDKAKRRNQEMQQKLLELTAENERLHKRVEQLSRDLSQVRHYFKQL
Dr-CEBPd_AAH56512 RHSPEYRQRRERNNIAVRKSRDKAKQRNLDMQQKMIELGAENERLHKTIDQLTRELSSLRNFFKQM
*Sp-CEBPa_XP_003725328 PGTHEYKQKRERNNIAVRKSREKTKTKNKELQDKVGELQEENTGLKKRVEGLAKELAVLRSSLTTR
Dm-CEBPL_SBC_NP_523843 KGTDEYRRRRERNNIAVRKSREKAKVRSREVEERVKSLLKEKDALIRQLGEMTNELQLHKQIYMQL
Aq-CEBPgL_XP_003382514 KCTDDYKDKRHRNNIAVRKSRSKFRKRVLETEKRVQELEENNAKLKNYVALLQKELAVLKGLFSSA
*Aq-CEBPaL_XP_011402720 KNSEEYRDKRERNNVAVRKSRNKMKMRAQETERRVHELEEENTALQNQVSLLLKELKVLKGLLSSA
Hs-CEBPg_AAC50201 RNSDEYRQRRERNNMAVKKSRLKSKQKAQDTLQRVNQLKEENERLEAKIKLLTKELSVLKDLFLEH
Mm-CEBPg_NP_034014 RNSDEYRQRRERNNMAVKKSRLKSKQKAQDTLQRVNQLKEENERLEAKIKLLTKELSVLKDLFLEH
Gg-CEBPg_NP_996741 RNSDEYRQRRERNNMAVKKSRLKSKQKAQDTLQRVTQLKEENERLEAKIKLLTKELSVLKDLFLEH
Xl-CEBPgL_XP_018116627 RGSEEYRQRRERNNMAVKKSRLKSKQKAQDTMQRVNQLKEENERLEAKIKLLTKELSVLKDLFLEH
Xl-CEBPgLH_NP_001089370 RGSEEYRQRRERNNMAVKKSRLKSKQKAQDTLQRVNQLKEENERLEAKIKLLTKELSVLKDLFLEH
Ac-CEBPg_XP_003227698 RASDEYRQRRERNNIAVKKSRLKSKQKAQDTLQRVNQLKEENERLEAKIKLLTKELSVLKDLFLEH
Dr-CEBPg_NP_571961 KDSDEYRQRRERNNLAVKKSRMRSKQKAQDTQQRVNELKEENERLEAKIKLLSKELSVLKDLFLEH
Sp-CEBPg_XP_003728584 SDSDEYKRRRERNNEAVRKSRQKSRQKASETEVRVTELKKENADLEQRVTLLHKELELLKDLFLTH
Ob-CEBPgL_XP_014783582 KSGDIYKKRRLRNNIAVRKCRSKNRMKAKETIDRVTRLRKENQDLQQKIQTLNKELSFLRDLFVGH
Ob-CEBPbL_XP_014769329 VTDKEYRNKRERNNVAVRKSRIKAKSKRVVTQNRVSQLKEENKQLEEKIKTLVQELKLCRSLFVQK
Hr-CEBPL_XP_009011126 KNNPNYIEKRERNNESVRKSRDKARQKQMETMEKVKVLTEENSRLNNKVNDLLKELEILKSLFANI
Ce-CEBPL_T15174 DDEDDYSTKRKRNNEAVNRTRQKKRQEENDTAEKVDELKKENETLERKVEQLQKELSFLKEMFMAY
*Mb-bZIP-TF5_XP_001743794 EDSNNYRIKRIRNNEAVRRCRIKKKQEMEEKAMRLELLEHKVSDLENCNRKLSELIVEQQKEIQRL
Co-ATF4L2_XP_004349044 DLPQRKLSRRERNNIAVRRCRDKNREKSLAAKSQCETVAQENANLRVRIHSLEQEVSYLKSMLLSQ
*Co-bZIP-TF1_XP_004363551 ETLADYLDKRKKNNDAVKKCRARKRMAVVATEEECQRLSGENASLRDRVGSLEAEVAYLKNLLISA
Ob-HP8-BRLZ1_XP_014769996 RKDQCYWEKRRKNNEAARRSREKRRLHDMALEKRIVELSRESCILRTQLYAVKKRYGIPREEPIIL
Hr-HP8L_XP_009021353 KKDKEYWMKRQKNNAAAKKSREKRRLNDVVLTNQIVQLTNENKRLKVELQAIKQRFGLSISSPY..
Ob-HP8-BRLZ2_XP_014769996 RLDPKYVERRKRNNEAARKCRENRKALTKLREVKSSYLESENGKLRNEMNDLQEEVKQLRELLEKK
Hs-CHOP_P35638 EDQGRTRKRKQSGHSPARAGKQRMKEKEQENERKVAQLAEENERLKQEIERLTREVEATRRALIDR
Mm-CHOP_AAH13718 EEQGRTRKRKQSGQCPARPGKQRMKEKEQENERKVAQLAEENERLKQEIERLTREVETTRRALIDR
Dr-CHOP_XP_005166228 PPVSSSRKRKRGGACSASPGKKSRREREQENERKVQELTDQNERLKAEIERLGEEVQRTRRALIER
L

fig. S13

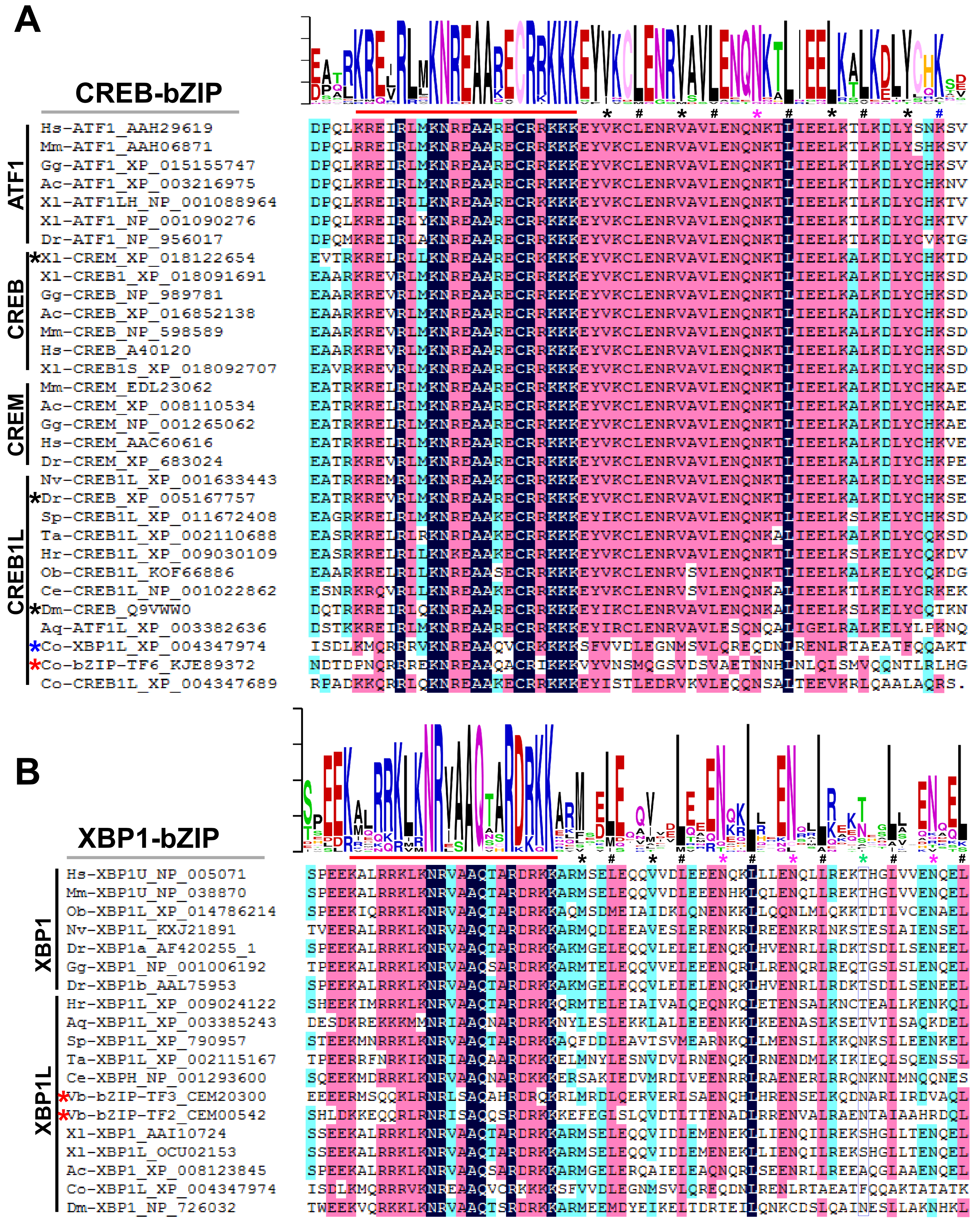
A
CREB-bZIP
ATF1
CREB
CREM
CREB1L
Hs-ATF1_AAH29619
Mm-ATF1_AAH06871
Gg-ATF1_XP_015155747
Ac-ATF1_XP_003216975
Xl-ATF1LH_NP_001088964
Xl-ATF1_NP_001090276
Dr-ATF1_NP_956017
*Xl-CREM_XP_018122654
Xl-CREB1_XP_018091691
Gg-CREB_NP_989781
Ac-CREB_XP_016852138
Mm-CREB_NP_598589
Hs-CREB_A40120
Xl-CREB1S_XP_018092707
Mm-CREM_EDL23062
Ac-CREM_XP_008110534
Gg-CREM_NP_001265062
Hs-CREM_AAC60616
Dr-CREM_XP_683024
Nv-CREB1L_XP_001633443
*Dr-CREB_XP_005167757
Sp-CREB1L_XP_011672408
Ta-CREB1L_XP_002110688
Hr-CREB1L_XP_009030109
Ob-CREB1L_KOF66886
Ce-CREB1L_NP_001022862
*Dm-CREB_Q9VWW0
Aq-ATF1L_XP_003382636
*Co-XBP1L_XP_004347974
*Co-bZIP-TF6_KJE89372
Co-CREB1L_XP_004347689
B
XBP1-bZIP
XBP1
XBP1L
Hs-XBP1U_NP_005071
Mm-XBP1U_NP_038870
Ob-XBP1L_XP_014786214
Nv-XBP1L_KXJ21891
Dr-XBP1a_AF420255_1
Gg-XBP1_NP_001006192
Dr-XBP1b_AAL75953
Hr-XBP1L_XP_009024122
Aq-XBP1L_XP_003385243
Sp-XBP1L_XP_790957
Ta-XBP1L_XP_002115167
Ce-XBPH_NP_001293600
*Vb-bZIP-TF3_CEM20300
*Vb-bZIP-TF2_CEM00542
Xl-XBP1_AAI10724
Xl-XBP1L_OCU02153
Ac-XBP1_XP_008123845
Co-XBP1L_XP_004347974
Dm-XBP1_NP_726032

**fig. S14**

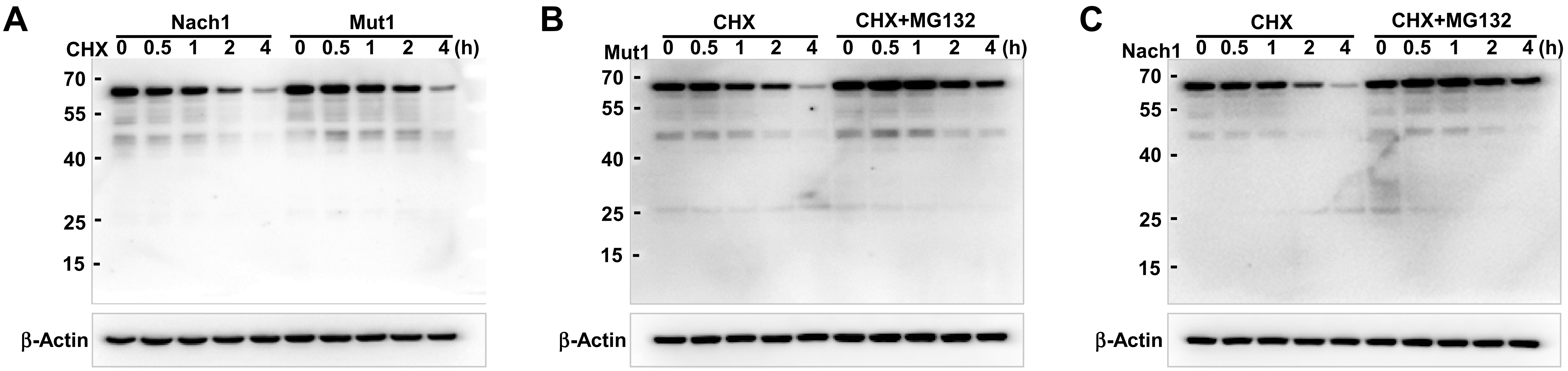
A
Nach1
Mut1
CHX 0 0.5 1 2 4 0 0.5 1 2 4 (h)
70
55
40
25
15
β-Actin
B
CHX
CHX+MG132
Mut1 0 0.5 1 2 4 0 0.5 1 2 4 (h)
70
55
40
25
15
β-Actin
C
CHX
CHX+MG132
Nach1 0 0.5 1 2 4 0 0.5 1 2 4 (h)
70
55
40
25
15
β-Actin
